# Quantitative analysis of the physiological contributions of glucose to the TCA cycle

**DOI:** 10.1101/840538

**Authors:** Shiyu Liu, Ziwei Dai, Daniel E. Cooper, David G. Kirsch, Jason W. Locasale

## Abstract

The carbon source for catabolism *in vivo* is a fundamental question in metabolic physiology. Limited by data and rigorous mathematical analysis, controversy exists over the nutritional sources for carbon in the tricarboxylic acid (TCA) cycle under physiological settings. Using isotope-labeling data *in vivo* across several experimental conditions, we construct multiple models of central carbon metabolism and develop methods based on metabolic flux analysis (MFA) to solve for the preferences of glucose, lactate, and other nutrients used in the TCA cycle across many tissues. We show that in nearly all circumstances, glucose contributes more than lactate as a nutrient source for the TCA cycle. This conclusion is verified in different animal strains from different studies, different administrations of 13C glucose, and is extended to multiple tissue types. Thus, this quantitative analysis of organismal metabolism defines the relative contributions of nutrient fluxes in physiology, provides a resource for analysis of *in vivo* isotope tracing data, and concludes that glucose is the major nutrient used for catabolism in mammals.

## INTRODUCTION

Cellular metabolism that resides within tissues utilizes many metabolites as their source in the TCA cycle such as glucose, lactate, amino acids, and fatty acids. As part of systemic metabolism, each cell has unique preferences for the utilization of particular metabolites, which is influenced by tissue type, cell state, environmental factors such as nutrition and physiological status. The nutrient preferences are critical for normal organ function, and closely linked to disease. For example, the fermentative glucose metabolism known as the Warburg effect has been widely found in numerous types of healthy and malignant cells (Liberti and Locasale, 2016), but glucose utilization is highly variable and depends on genetics and environment (Faubert et al., 2017; Feron, 2009; Hensley et al., 2016). Those specific metabolic fluxes could be potential targets for cancer treatment (Liberti et al., 2017; Sonveaux et al., 2008). For other tissues like the myocardium, the energy contribution from fatty acids, glucose, lactate and others are thought to directly reflect its nutrient and oxygen availability, and have important roles in cardiology (Kodde et al., 2007; Ma et al., 2019). Therefore, an investigation of nutrient source utilization in physiological conditions is of utmost importance.

To quantitate different nutrient sources, isotope-labeling-based methods have long been used. Cells or animals are fed or infused with isotopically-labeled substrates, and labeling ratios of metabolites are analyzed by mass spectrometry (MS) or nuclear magnetic resonance (NMR). Previous studies have used these data to qualitatively explain the contribution of nutrient sources to the TCA cycle (Stanley et al., 1988). However, those studies have been limited by measurements that often included only a few metabolites. Recent studies have looked to quantitatively measure the utilization of nutrient sources at the systemic level using metabolic flux analysis (MFA) (Hui et al., 2017; Jang et al., 2019; Neinast et al., 2019). MFA is a mathematical framework that seeks a solution of metabolic fluxes that best fits the isotope labeling data (i.e. using machine learning or artificial intelligence) for a given biochemical reaction network (Dai and Locasale, 2017; Zamboni et al., 2009). The biochemical model used is essential for the resulting solutions. For instance, reversible (i.e. exchange) fluxes of metabolites between tissue and plasma are almost always significant and may highly influence isotope labeling patterns (Witney et al., 2011). However, many MFA models do not consider exchange fluxes (Hui et al., 2017). Another important point is the heterogeneity of metabolism. Some studies have shown that metabolic heterogeneity exists widely in within and between lung cancers (Hensley et al., 2016). Organismal metabolism relies on mutual cooperation between tens of organs and tissues. However, most current MFA models consider the flux calculation in one kind of tissue and assume the tissue is a homogenous system.

To investigate the quantitative selection of nutrient sources of entry into the TCA cycle under physiological conditions, we developed a framework to overcome current challenges. Multiple tissues are considered, linked by circulation. This model also uses the MFA framework and requires isotope-labeling data for different tissues to fit fluxes in different compartments. Surprisingly we found that under physiological conditions, as we validated using different animal models and experimental isotope labeling conditions, most tissues utilize circulating glucose more than lactate for the TCA cycle which may challenge current dogma in metabolic physiology.

## RESULTS

### Model construction and flux analysis

In the fasting state, systemic metabolism involves a source tissue (usually liver) that converts circulating lactate to glucose in blood, and a sink tissue that consumes glucose back to lactate, which is referred as the Cori cycle (Nelson et al., 2017) (Figure 1A). Glucose and lactate in the source and sink tissues are interconverted through pyruvate. Sink and source tissues are connected through plasma, which allows for the transport of glucose and lactate (Figure 1B).

**Figure 1.**
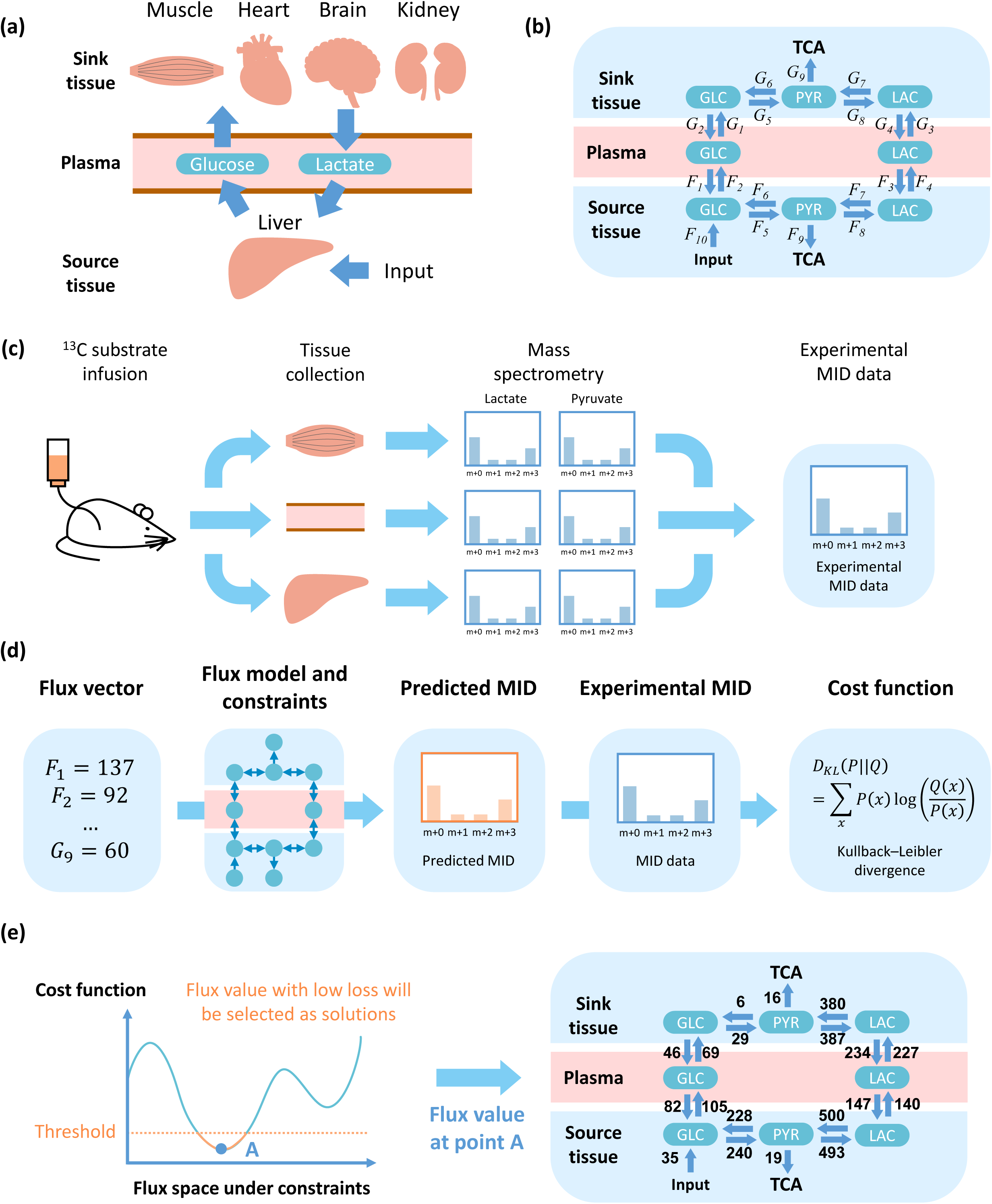
General methodology and flux analysis. (a) Diagram of metabolite exchange between source and sink tissues. Glycogen, amino acids and other nutrition source are utilized to supplement glucose in the source tissue. (b) Three components (source tissue, plasma and sink tissue) and two circulating metabolites (lactate and glucose). (c) Data acquisition. Tissues of ^13^C-infused mice are extracted and analyzed by mass spectrometry. Distribution of mass isotopomers for metabolites, such as glucose, lactate and pyruvate, are used to solve for the fluxes (b). (d) Definition of cost function. The flux vector is used to predict MID of target metabolites, and compared with experimental MID to calculate cost function. (e) Schematic and example of a feasible solution. The solution with cost function lower than a threshold is considered as feasible solution and will be utilized in the following analysis.

Fluxes are computed based on data from mass spectrometry (MS) in ^13^C-glucose infused mice as follows. After infusion, tissues are collected and analyzed by MS. Metabolites with ^13^C at different positions are distinguished and their relative abundance is referred to as the mass isotopomer distribution (MID) (Figure 1C). MID data are then used to fit the fluxes in the model. Given a set of fluxes, MIDs are calculated and compared with experimental data. The difference (i.e. cost function) between the estimated MIDs and experiments, measured by a standard metric used in Information Theory, the Kullback-Leibler divergence (Kullback and Leibler, 1951), is minimized to find a set of fluxes that best fits the data. Next, statistical sampling is conducted to find all sets of fluxes that can be considered as valid solutions (Figures 1D, E, supplementary information). Additional constraints are then introduced to ensure the simulated fluxes are physiological feasible, such as requirements for minimal TCA flux values in the source and sink tissues (supplementary information).

The model was first fit and fluxes were computed using data from a recent study (Hui et al., 2017). Among all calculations of fluxes obtained from our algorithmic procedure (Figures 1C-E), the MIDs of most metabolites can be predicted by the current model (Figure S1A-H), and the values of the fluxes in the model are physiologically feasible (Figure S1I). The value of the cost function for the set of fluxes computed is also significantly lower that what is obtained from considering randomized data indicating that the values of fluxes computed are statistically significant (methods, Figure S1J-P).

### Glucose contributions in different tissues

The flux network can be mathematically defined with a simplified diagram: the TCA cycle in the source and sink tissue is fed by two fluxes from glucose and lactate in plasma (Figure 2A, supplementary information). Non-negative contribution fluxes to TCA cycle from glucose (*F*_*glc*_ in source tissue and *G*_*glc*_ in sink tissue) or from lactate (*F*_*lac*_ in source tissue and *G*_*lac*_ in sink tissue) are calculated from net fluxes of related reactions and diffusion (supplementary information). From the computed fluxes, two glucose contribution ratios, a local one *R*_*glc*_ and a global one 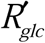, are defined to reflect the relative ratio of glucose contribution to the TCA cycle in sink tissue only or in complete model, which includes both sink and source tissue, respectively. If *R*_*glc*_ or 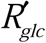 is higher than 0.5, it implies that glucose contributes more than lactate to TCA cycle. On the contrary, if it is lower than 0.5, lactate contributes more than glucose (Figure 2B, S2A). The global ratio 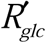 reflects the glucose contribution in the complete model, while the local value *R*_*glc*_ distinguishes the glucose flux from the source tissue and from circulating metabolites into the sink tissue.

**Figure 2.**
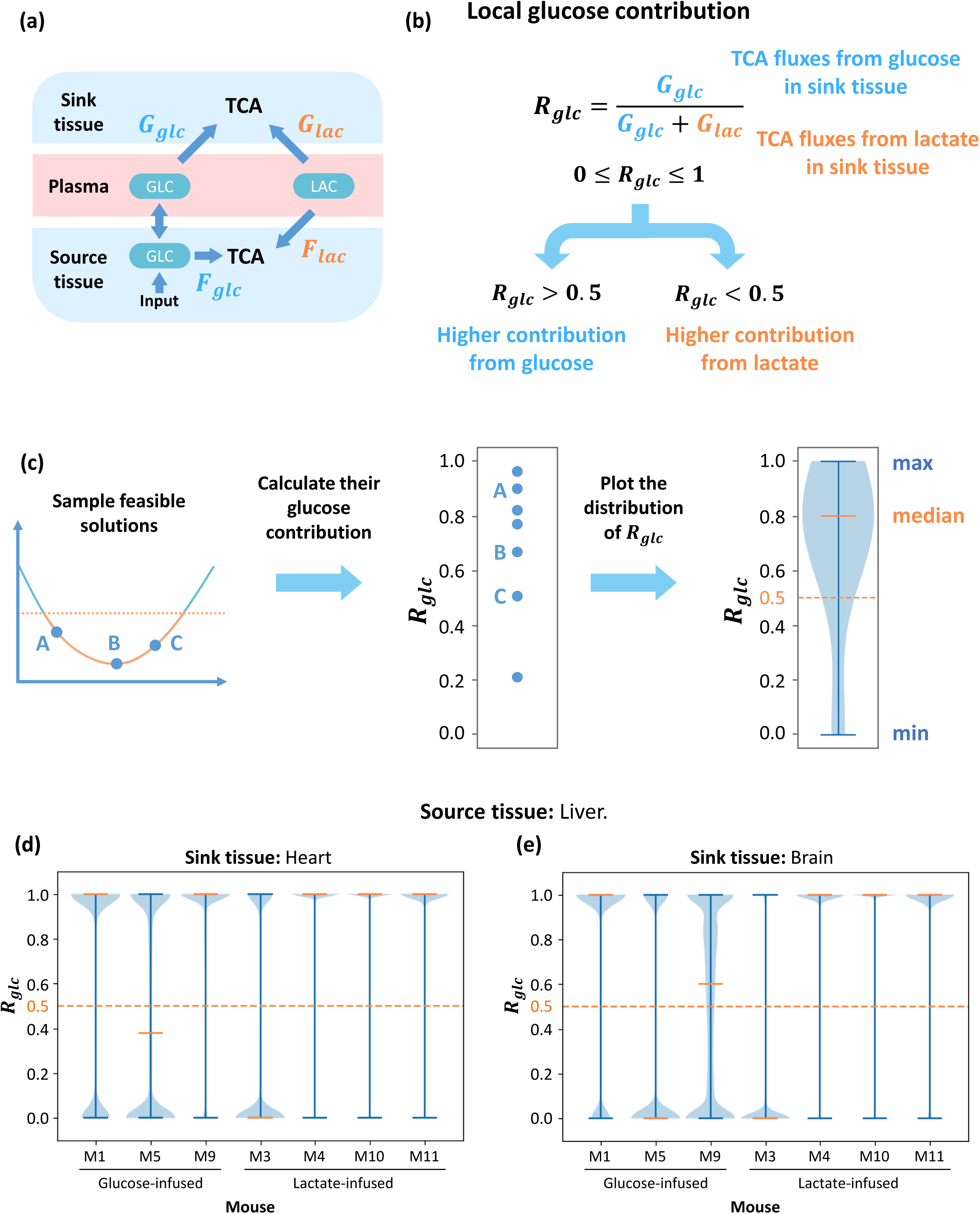
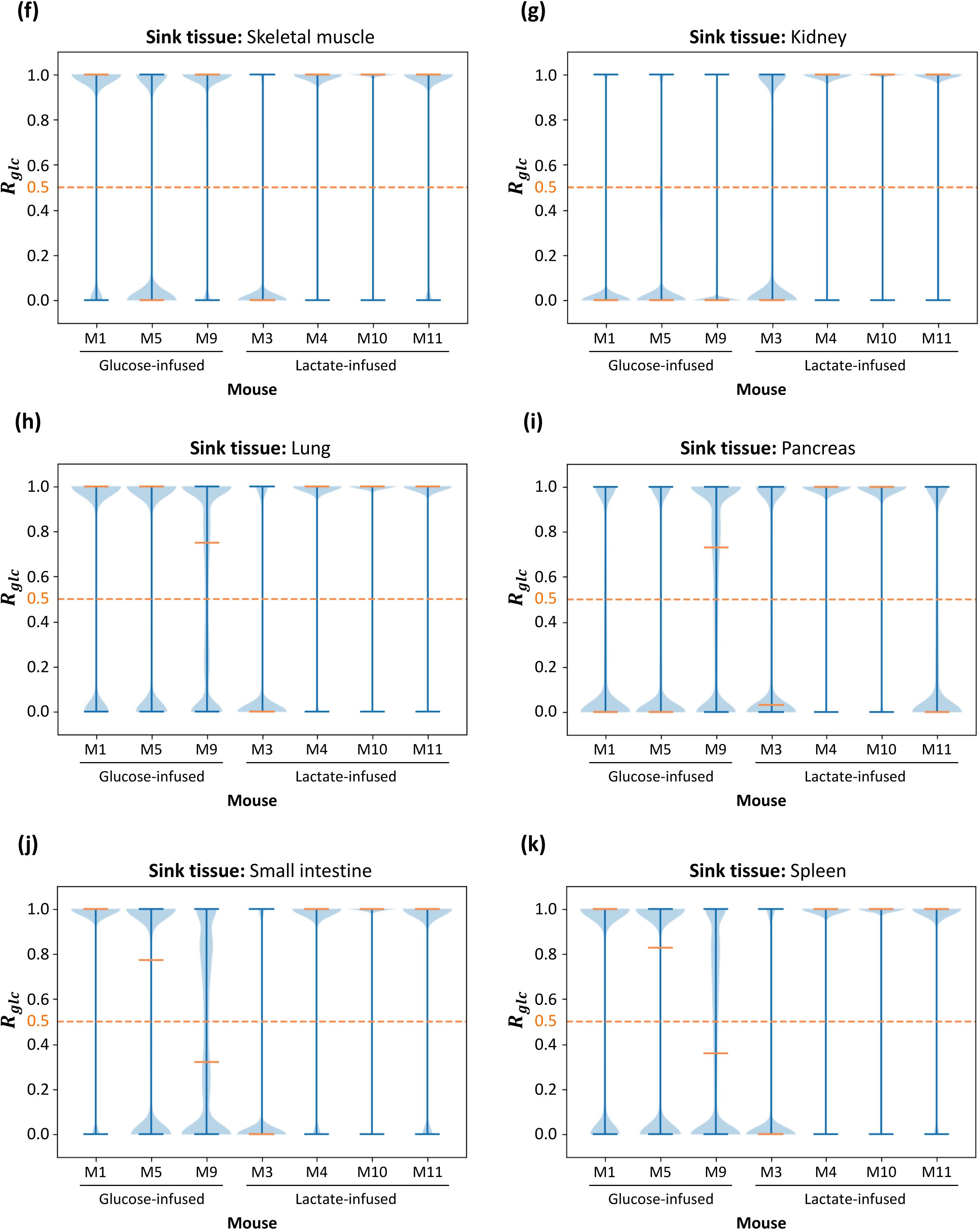
Contribution to the TCA cycle from circulating glucose. (a) Diagram of contribution fluxes. Glucose and lactate contribute to the TCA cycle by *F*_*glc*_ and *F*_*lac*_ in the source tissue, while *G*_*glc*_ and *G*_*lac*_ are related to the sink tissue. The direction of net flux between circulating glucose and glucose in source tissue is variable in different solutions. (b) Definition of global glucose contribution ratio *R*_*glc*_ based on fluxes in (a). The global glucose contribution *R*_*glc*_ is defined as the relative ratio of glucose contribution flux to total contribution flux in sink tissue. *R*_*glc*_ is a scalar between 0 and 1, and higher *R*_*glc*_ represents higher glucose contribution to the TCA cycle. (c) Procedure to compute distribution of glucose contribution. Feasible solutions are sampled and glucose contribution ratios are calculated. The distribution of glucose contribution is displayed by a violin plot. (d-k) Distribution of local glucose contribution based on models with different sink tissues. For each sink tissue, the source tissue is liver, and contribution ratio is calculated from data in 7 different mice. For most kinds of sink tissue, the median of glucose contribution is higher than 0.5 in most mice, which means glucose contributes more than lactate to the TCA cycle. The orange dash line represents 0.5 threshold. Data set is from glucose-infused mice (M1, M5, M9) and lactate-infused mice (M3, M4, M10, M11) in Hui et al, 2017.

To evaluate the glucose contribution, feasible solutions are sampled from the solution space and displayed in a violin plot (Figure 2C). The source tissue is the liver and the sink tissues are set as heart, brain, skeletal muscle, kidney, lung, pancreas, small intestine and spleen. For each combination of sink and source tissue, the model is fit with data from mice infused by glucose and lactate. The local glucose contribution ratios *R*_*glc*_ have a bimodal distribution (supplementary information), and they concentrate around 1 in models for almost all sink tissues in most of infused mice (Figure 2D-K). For the global glucose contribution ratios 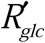, the median of the distribution in all kinds of sink tissue is higher than 0.5 (Figure S2B-I). Therefore, those results show that in almost all cases glucose contributes more than lactate does to the TCA cycle.

The results from these two-tissue models rely on MID data and some parameters. To evaluate these dependencies, we implemented a Monte Carlo based sensitivity analysis (Shestov et al., 2014). First, original data and parameters are perturbed randomly. The perturbed values are used to calculate distribution of the local contribution *R*_*glc*_ as previously described. After this process, the median of this distribution under each individual perturbation is collected, and the distribution of median *R*_*glc*_ reflects its sensitivity to data and parameters (Figure 3A). Results show that the median value of *R*_*glc*_ is very robust to perturbations in glucose circulatory flux and input flux, but more sensitive to the value of lactate circulatory flux and the MID data (Figure 3B-E). However, in most parameter sets, the median *R*_*glc*_ is still higher than 0.5 (Figure 3C, 3E). These results demonstrate the robustness of the conclusion that glucose contributes more than lactate to the TCA cycle under physiological conditions.

**Figure 3.**
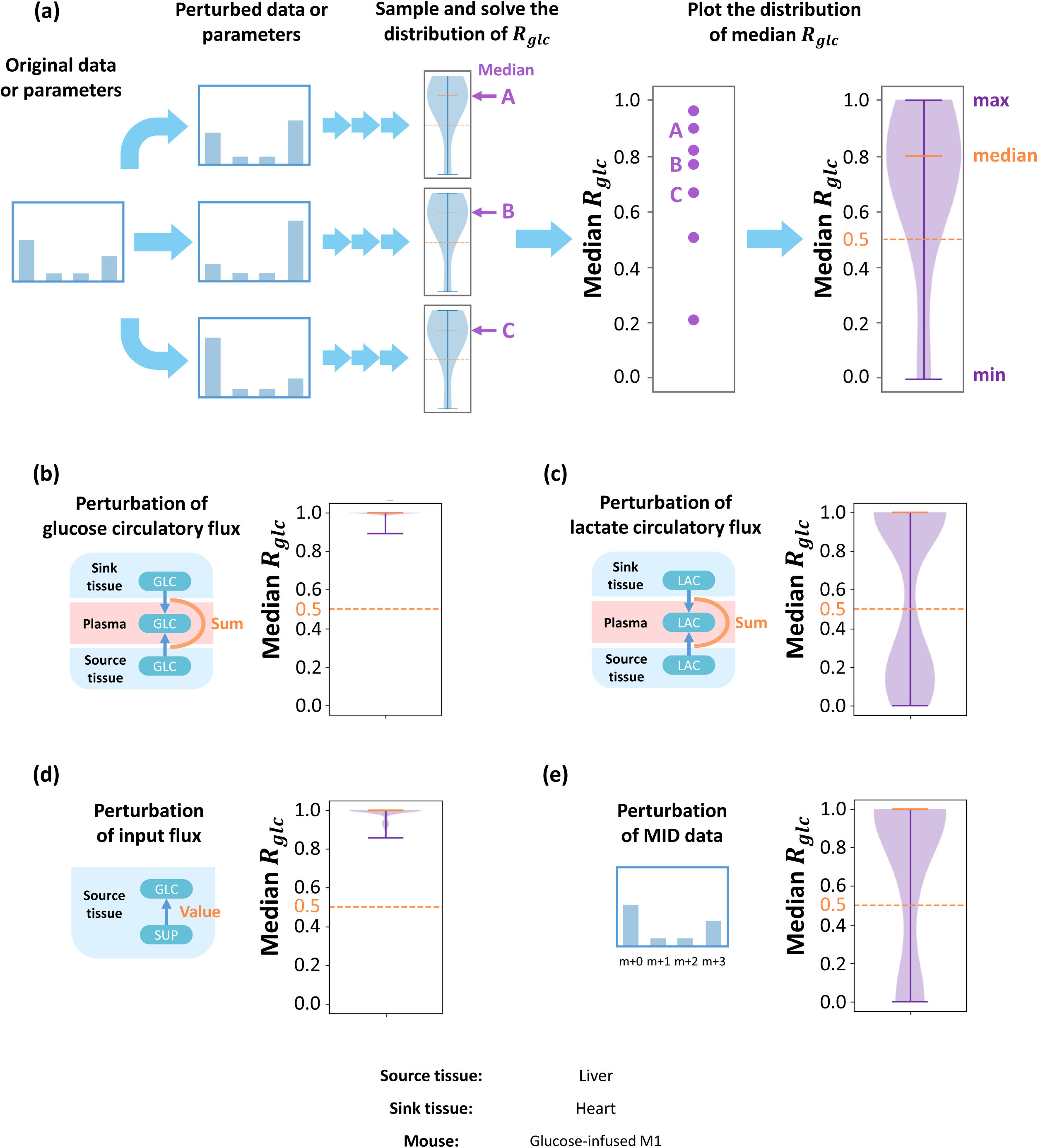
Parameter sensitivity analysis. (a) Original MID data or constraint parameters are randomly perturbed and used in the following analysis. The resulting distribution of the local glucose contribution for each perturbation is calculated, and their medians are collected. Distribution of medians reflects parameter sensitivities for this model. The distribution of medians under perturbation of glucose circulatory flux (b), lactate circulatory flux (c), input flux in source tissue (d) and MID data (e). Although the local contribution ratio is more sensitive to lactate circulatory flux and MID data, most of the medians are above the 0.5 threshold, which implies that under most perturbations, glucose contributes more than lactate to the TCA cycle. Data set is from glucose-infused mouse M1 in Hui et al, 2017. Source tissue is liver and sink tissue is heart.

One confounding issue is that the process of tissue harvesting may induce ischemia and hypoxia. Hypoxia will induce elevated glycogenolysis in source tissue and glycolysis in sink tissue, which may significantly change measured MID of metabolites (Figure S3A). To estimate its effect on the final conclusion, a correction is introduced to simulate these effects under hypoxia. Measured MIDs of glucose in source tissue and lactate in sink tissue are assumed to be a mixture of 80% real MID in physiological state, and 20% MID of newly synthesized metabolites in elevated reactions under hypoxia (Figure S3B). Specifically, glucose in source tissue is assumed to be mixed with unlabeled glucose, and lactate in sink tissue is assumed to be mixed with lactate synthesized from pyruvate, which has same MID as pyruvate. Therefore, the physiological MID can be solved for and utilized for the same analysis of glucose contribution. Compared to results before the correction, conclusions were not altered, and in most cases glucose contributes more than lactate is robust to hypoxia considerations (Figure S3C-D).

### Generality of the glucose contribution to the TCA cycle

To further investigate the generality of this conclusion, we considered a different animal strain, different diet and different infusion protocol with mice infused with ^13^C-glucose at a higher infusion rate which is one of the key technical variables of consideration in these studies (Ayala et al., 2010). In addition to our analysis of published data (Hui et al., 2017), these new experiments expand the scope of physiological variables (Figure 4A). These data are referred to as “high-infusion rate”, while the previous analysis is referred to as “low-infusion rate”. Importantly, with the higher infusion rate, the glucose, lactate and insulin levels in plasma are not significantly altered during the infusion (Figure 4B). Because of the higher infusion rate, an input flux *J*_*in*_ in plasma is added to the model to capture the infusion operation (Figure 4C). The amount of ^13^C labeling increases with the infusion rate, and with a higher infusion rate, the model predicts the MIDs (Figure S4A). In this case, the cost function is also significantly lower than that obtained from a random unfitted control for the 4 glucose-infused mice in the higher infusion rate experiments (Figure 4D), and the value of all fluxes are physiologically feasible (Figure S4B).

**Figure 4.**
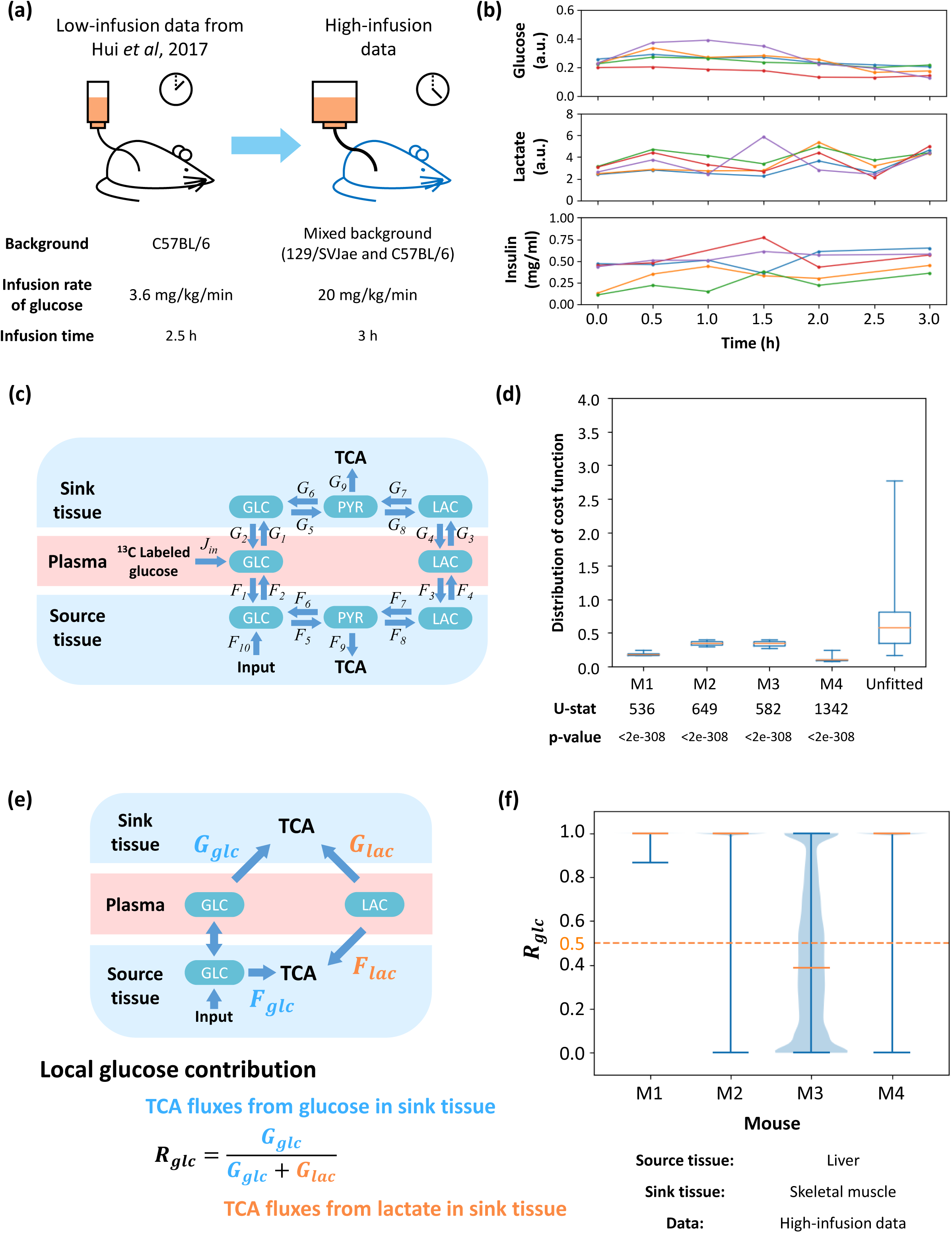
Robustness of results regarding animal strain and infusion rate. (a) Diagram of comparison between two experiments. A higher infusion rate and longer infusion time are introduced, which leads to higher abundance of ^13^C labeling in most metabolites. The genetic background and diet are also different from previous experiments. (b) Time-course data for concentrations of glucose, lactate and insulin in plasma during infusion. Each color represents a specific mouse. In the insulin measurement, a data point at 1h of red line is removed because of a significantly abnormal value. (c) Structure of high-infusion model. The main difference is ^13^C labeled infusion to glucose in plasma. (d) Distribution of cost function fitted with data from different mice or unfitted control data. U-statistics of a rank-sum test and p-values are displayed. (e) Definition of local glucose contribution *R*_*glc*_. Glucose and lactate in plasma contribute to the TCA cycle in the source and sink tissue. Direction of net flux between circulating glucose and glucose in source tissue is variable in different solutions. The local glucose contribution *R*_*glc*_ is defined as the relative ratio of glucose contribution flux to total contribution flux in sink tissue. (f) Distribution of local glucose contribution shows glucose contributes more than lactate to the TCA cycle in most cases. Fits from different mice are displayed. In all subfigures, the source tissue is liver and sink tissue is skeletal muscle.

As defined previously, the local contribution *R*_*glc*_ and global contribution 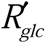 to the TCA cycle are calculated for the pair of source tissue (liver) and sink tissue (skeletal muscle) for all mice (Figure 4E, S4C). The analysis shows that in most mice, *R*_*glc*_ and 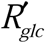 are both higher than 0.5, again implying that glucose contributes more than lactate to the TCA cycle (Figure 4F, S4D).

### Glucose contribution upon consideration of multiple tissue interactions

The current model is based on source and sink tissues. However, mammals consist of tens of different tissues which cooperate and interact. To demonstrate the utility of this model to multiple tissue compartments, more sink tissues are introduced and the glucose contribution under these conditions are analyzed. This model contains one source tissue and two sink tissues, which are connected by glucose and lactate in plasma (Figure 5A, 5B). This model is fit with the low-infusion rate data, in which source tissue is liver and two sink tissues are combinations from heart, brain and skeletal muscle. The fitting is sufficiently precise (Figure S5A), implying that computed fluxes are physiologically feasible (Figure S5B). The cost functions of all combinations are also significantly lower than a random unfitted control (Figure S5C). In this model, glucose and lactate in plasma can contribute to the TCA cycle through three kinds of tissue, and therefore the definitions of local and global glucose contribution ratios *R*_*glc*_ and 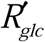 are slightly modified (Figure 5C, S5D). Fitting results show in all three combinations of two sink tissues, glucose contributes more than does lactate to the TCA cycle regardless of the definition of glucose contribution ratio (i.e. local or global contribution ratio) used (Figure 5D, S5E).

**Figure 5.**
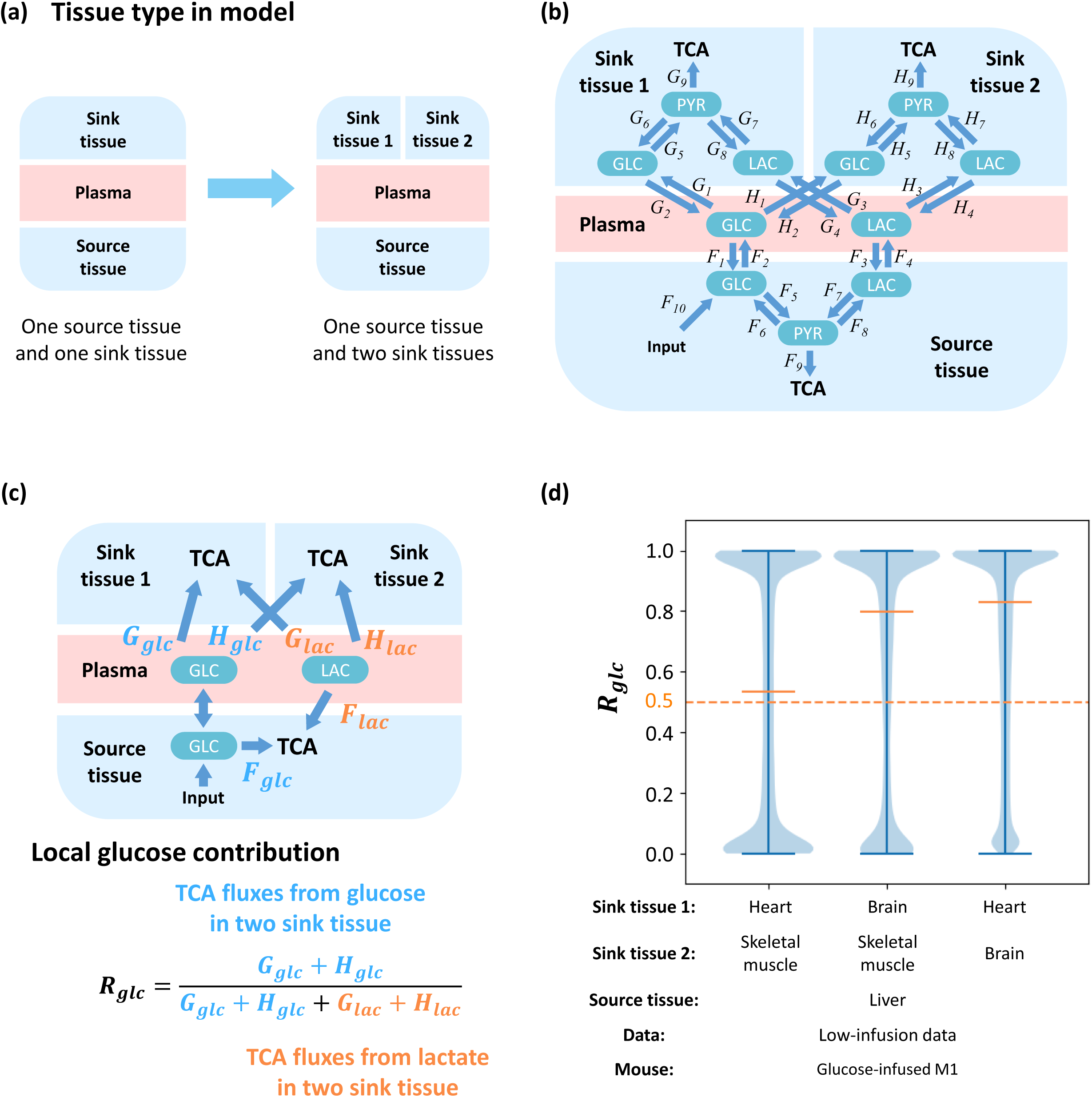
Flux analysis across multiple tissues. (a) A model with additional sink tissues. (b) Structure of the multi-tissue model. One source tissue and two sink tissues are connected by glucose and lactate in the plasma. (c) Definition of local glucose contribution *R*_*glc*_. Glucose and lactate can contribute to TCA by *F*_*glc*_ and *F*_*lac*_ in the source tissue, *G*_*glc*_ and *G*_*lac*_ in the sink tissue 1, and *H*_*glc*_ and *H*_*lac*_ in sink tissue 2, respectively. Direction of net flux between circulating glucose and glucose in source tissue is variable in different solutions. The local glucose contribution is defined as the relative ratio of glucose contribution flux to total contribution flux in two kinds of sink tissue. (d) Distribution of local glucose contribution shows glucose contributes more than lactate to the TCA cycle in all combinations of sink tissues. The model is fit by glucose-infused mouse M1 from the low-infusion data in Hui et al, 2017. The source tissue is liver and the sink tissue 1 and 2 are two from heart, brain and skeletal muscle respectively.

### Glucose contribution upon consideration of multiple nutrient sources

The current analysis considers two circulating metabolites as sources for the TCA cycle: glucose and lactate. However, many other metabolites circulate and are exchanged between tissue and plasma, such as acetate, alanine and pyruvate (Hui et al., 2017; Liu et al., 2018). Therefore, to investigate the applicability of this model, circulating pyruvate is introduced (Figure 6A). Circulating pyruvate can also represent other nutrient sources including but not limited to alanine, glutamine, acetate, or fatty acids. In this model, circulating pyruvate is not only exchanged with sink and source tissue, but also converted to lactate in plasma. Glucose and lactate in plasma can also be directly converted to pyruvate (Figure 6B). This model predicts the experimental MID with both low-infusion rate and high-infusion rate data (Figure S6A, S6B) with the physiologically feasible fluxes (Figure S5C, S5D). The distribution of values of the cost function is also significantly lower than random unfitted control in all kinds of sink tissue fitted with the low-infusion data (Figure S6E), or in the skeletal muscle fitted with the high-infusion data indicating statistical significance (Figure S5F).

**Figure 6.**
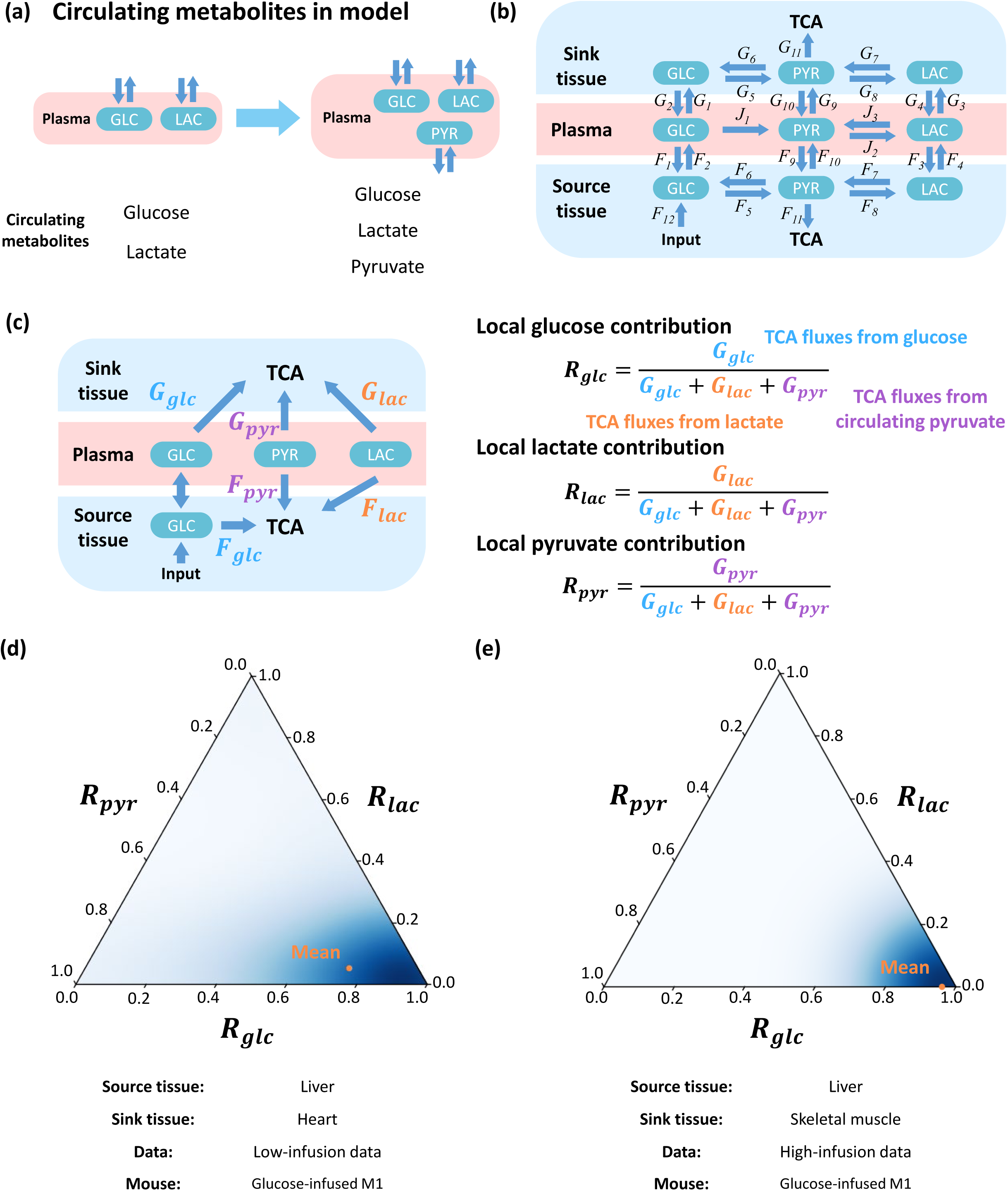
Model with multiple circulating metabolites feeding the TCA cycle. (a) Incorporation of additional circulating metabolites. (b) The structure of the model. The source tissue and sink tissue are connected with glucose, lactate and pyruvate in the plasma. (c) Definition of local contribution from metabolites *R*_*glc*_, *R*_*lac*_ and *R*_*pyr*_. Glucose, lactate and pyruvate can contribute to the TCA cycle by *F*_*glc*_, *F*_*lac*_ and *F*_*pyr*_ in source tissue, and *G*_*glc*_, *G*_*lac*_ and *G*_*pyr*_ in the sink tissue. Direction of net flux between circulating glucose and glucose in source tissue is variable in different solutions. The local contribution ratios of three metabolites *R*_*glc*_, *R*_*lac*_ and *R*_*pyr*_ are defined by the relative ratio of the contribution flux from each metabolite to total contribution flux of all three metabolites in sink tissue. (d) Ternary plot to display distributions of local contributions from three metabolites. The orange point indicates average level. The model is fit by glucose-infused mouse M1 from low-infusion data. The source tissue is liver and sink tissue is heart. (e) Analysis and results as in (d) but for additional high-infusion system of different animal strain, different diet and different infusion protocol. The model is fitted by glucose-infused mouse M1 from the high-infusion data. The source tissue is liver and sink tissue is skeletal muscle.

Because circulating glucose, lactate and pyruvate each contributes to the TCA cycle in source and sink tissues, the local contribution ratios of the three metabolites *R*_*glc*_, *R*_*lac*_ and *R*_*pyr*_ need to be calculated individually, as well as three global contribution ratios 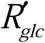, 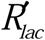 and 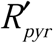, and the sum of the three local or global contribution ratios equals to 1 (Figure 6C, S6H). The distribution of three ratios can be displayed by a ternary plot (Marc et al., 2019, supplementary information). For the low-infusion rate data, the local contribution from glucose is predominantly higher than lactate and pyruvate (Figure 6D), and the conclusion is similar when the sink tissue in the model is replaced by other types of tissue (Figure S6G). For the global contribution, contribution from glucose is close to or slightly lower than lactate, which are both significantly higher than pyruvate (Figure S6I). The situation is similar in the high-infusion rate data, in which the local contribution from glucose markedly dominates in all sampled solutions, but the global contribution from glucose is closed to lactate (Figure 6E. S6J). Therefore, in a model with more metabolites in circulatory system, circulating glucose contributes more than lactate to the TCA cycle in all kinds of sink tissue, and has similar contribution than does lactate.

## DISCUSSION

The nutrient sources for the TCA cycle have long been of interest. However, due to difficulties in data acquisition and mathematical analysis, quantitative studies under physiological conditions are still rare. With advances in mass spectrometry and mathematical modeling, *in vivo* flux analysis studies with isotope-labeling data have become a mainstay in the study of metabolic physiology. Previous studies have measured TCA cycle source utilization by MFA. However, with the development of these new mathematical tools, our study challenges some key conclusions that form the current consensus for the relative contributions of lactate and glucose to the TCA cycle. For example, it was reported that lactate is the major energy source for most tissues and tumors (Hui et al., 2017; Jin et al., 2019). Our results show that most organs uptake more glucose than lactate to fuel the TCA cycle. This conclusion also holds under various parameters, experimental conditions such as animal strain and diet, tissue type, tissue interactions and source metabolite number, which together indicate the robustness and generalizability of the conclusions. Our results, however, are consistent with conventional knowledge that glucose behaves as a primary energy source in cells and tissues, especially for neural systems (Nelson et al., 2017). Nevertheless, our results confirm that lactate is highly exchanged between tissue and plasma, while glucose is transferred from the liver to other organs. These phenomena appear to also be observed in recent studies on flux measurements in pigs (Jang et al., 2019).

In addition to addressing an important issue in metabolic physiology, our study provides a framework for metabolic flux analysis in physiological conditions. Compared with previous studies, the first improvement is that fluxes calculated by our model can capture more aspects of metabolic biochemistry. For example, the flux from pyruvate to glucose (G6/H6), the gluconeogenesis flux in sink tissue, relies on Phosphoenolpyruvate Carboxykinase (PEPCK), which only expresses in few kinds of tissue such as liver, kidney and adipose tissue (Geiger et al., 2013). Therefore, G6/H6 fluxes are very small in most of our fitting results (Figure S1I, S4B, S5B, S6C-D). Another example is high diffusion and exchange of lactate between tissue and plasma, which is usually overlooked, but captured by our model (F3/F4, G3/G4, H3/H4 in Figure S1I, S4B, S5B, S6C-D) and validated by experimental measurements (Jang et al., 2019). The second improvement is that, rather than fitting the model with a single solution, we sampled the entire high-dimensional solution space and analyzed all feasible results. Those millions of sampled points can cover more regions in solution space and precisely reflect real distribution of fluxes, especially in a complicated model. The third improvement is more complete analysis for parameter sensitivity than previous studies. This study verified the robustness of the conclusions not only under random perturbation of parameters and MID data, which accounts for uncertainties in experimental precision, but also may account for hypoxia which introduces systematic experimental bias. These analyses serve to extend much of the Metabolic Flux Analysis framework that was developed for cell systems to physiological conditions.

Another intriguing feature of this model is its generalizability and scalability. From a basic two-tissue version, this model is readily extended to compute fluxes from isotope patterns with higher infusion rates, more tissue types and more nutrient sources which could be useful to study for example different nutritional situations and pathophysiology states such as metabolic syndromes, diabetes and cardiovascular disease. The generality of this model allows for a broader usage in future research. More kinds of tissue can be introduced to better mimic the physiological condition such as the interaction between cancer and host organs. As the number of tissues increases, their roles could be more complicated rather than simple source and sink. For example, previous research indicates that the kidney may also have a significant contribution to net production of glucose in pigs (Jang et al., 2019). Second, more nutrient sources could be introduced and the metabolic network in each cell could also be expanded. The current model includes three nodes: glucose, pyruvate and lactate which capture fluxes in central carbon metabolism but could be extended into intermediary metabolism. Although sufficient for analyzing the contribution of macronutrients, studies of fatty acids, ketosis and amino acid metabolism will require a larger network. Nevertheless, the methodology contained within this model could be extended. For instance, subcellular compartmentalized metabolic flux analysis is also important (Lee et al., 2019). However, its application is usually restricted to the mitochondria and nucleus because of the difficulty in acquiring isotope-labeling data in each cellular compartment. On the other hand, interactions within heterogenous tissues could also be described by this model. It has been widely shown that cells in a tumor may express different metabolic states, and will compete or cooperate for many resources (Hensley et al., 2016). Quantitative methods based on this model may help to better describe those precise and complicated interactions.

Our ability to resolve metabolic fluxes is limited by the availability of high-quality data. Thus, limited by data and computational resources, this model only covers a small portion of biochemical reactions. Specifically, this model combines all fluxes in the TCA cycle into one unidirectional flux, because adding those fluxes and metabolites to the model will not largely improve fitting precision of current fluxes, but will increase the dimension of solution space and thus increase uncertainty of results (supplementary information). Therefore, this model may not perfectly fit MID of some metabolites connected with TCA. For example, pyruvate can feed the TCA cycle and change the MID of metabolites in it, but it can also be fed by cataplerotic fluxes of TCA cycle. Consequently, the MID of pyruvate will be coupled with metabolites in TCA cycle, and cannot be precisely described as the model currently stands. Another limitation is the high dimensionality of the solution space in light of limited available constraints. In our models, high dimensionality of the solution space requires sampling algorithms to measure the solution space. As the model expands, these algorithmic challenges become more difficult. Thus, more constraints must be introduced to reduce the dimensionality of the feasible solution space. For example, our study includes constraints from circulatory fluxes (Hui et al., 2017), and some MFA model uses fixed biomass fluxes as boundary conditions (Reid et al., 2018). However, the precision and generalizability of these external constraints have not been rigorously validated, and they may introduce bias. Comprehensive and precise model analysis requires more effort to establish reliable constraints as well as acquisition of metabolite data with more coverage and higher resolution.

## METHODS

### Data Sources

This study is based on two data sources: low-infusion data were obtained from infused fasting mice in previous work (Hui et al., 2017), while the high-infusion data were acquired based on following protocols.

### Reagents

Unless otherwise specified, all reagents were purchased from Sigma-Aldrich. Jugular vein catheters, vascular access buttons, and infusion equipment were purchased from Instech Laboratories. Stable isotope glucose were purchased from Cambridge Isotope Laboratories.

### Animal Models

All animal procedures were approved by the Institutional Animal Care and Use Committee (IACUC) at Duke University. Mouse models is from 8 to 10-week old, male and female mixed background (129/SVJae and C57BL/6) with a combination of alleles that have been previously described: Pax7^CreER-T2^, p53^FL/FL^, LSL-Nras^G12D^ and ROSA26^mTmG^ (Zhang et al., 2015). Mice were fed standard laboratory chow diets *ad libitum*.

### *In vivo*^13^C glucose infusions

To perform in vivo nutrient infusions, chronic indwelling catheters were placed into the right jugular veins of mice and animals were allowed to recover for 3-4 days prior to infusions. Mice were fasted for 6 hours and infused with [U-13C]glucose for 3 hours at a rate of 20 mg/kg/min (150 µL/hr). Blood was collected via the tail vein at 3 h and serum was collected by centrifuging blood at 3000g for 15 min at 4°C. At the end of infusions, tissues were snap frozen in liquid nitrogen and stored at -80°C for further analyses.

### Insulin measurement

The concentration of insulin in plasma is measured by Ultra Sensitive Mouse Insulin ELISA Kit from Crystal Chem.

### Metabolite extraction from tissue

Briefly, the tissue sample was first homogenized in liquid nitrogen and then 5 to 10 mg was weighed in a new Eppendorf tube. Ice cold extraction solvent (250 μl) was added to tissue sample, and a pellet mixer was used to further break down the tissue chunk and form an even suspension, followed by addition of 250 μl to rinse the pellet mixer. After incubation on ice for an additional 10 min, the tissue extract was centrifuged at a speed of 20 000 g at 4 °C for 10 min. 5 μl of the supernatant was saved in -80 °C freezer until ready for further derivatization, and the rest of the supernatant was transferred to a new Eppendorf tube and dried in a speed vacuum concentrator. The dry pellets were reconstituted into 30 μl (per 3 mg tissue) sample solvent (water:methanol:acetonitrile, 2:1:1, v/v) and 3 μl was injected to LC-HRMS.

### HPLC method

Ultimate 3000 UHPLC (Dionex) was used for metabolite separation and detection. For polar metabolite analysis, a hydrophilic interaction chromatography method (HILIC) with an Xbridge amide column (100 × 2.1 mm i.d., 3.5 µm; Waters) was used for compound separation at room temperature. The mobile phase and gradient information were described previously. 2-hydrazinoquinoline derivatives were measured using reversed phase LC method, which employed an Acclaim RSLC 120 C8 reversed phase column (150 × 2.1 mm i.d., 2.2 μm; Dionex) with mobile phase A: water with 0.5% formic acid, and mobile phase B: acetonitrile. Linear gradient was: 0 min, 2% B; 3 min, 2% B; 8 min, 85% B;9.5 min, 98% B; 10.8 min, 98% B, and 11 min, 2% B. Flow rate: 0.2 ml/min. Column temperature: 25 °C.

### Mass Spectrometry

The Q Exactive Plus mass spectrometer (HRMS) was equipped with a HESI probe, and the relevant parameters were as listed: heater temperature, 120 °C; sheath gas, 30; auxiliary gas, 10; sweep gas, 3; spray voltage, 3.6 kV for positive mode and 2.5 kV for negative mode. Capillary temperature was set at 320°C, and S-lens was 55. A full scan range was set at 70 to 900 (*m/z*) with positive/negative switching when coupled with the HILIC method, or 170 to 800 (*m/z*) at positive mode when coupled with reversed phase LC method. The resolution was set at 140 000 (at *m/z* 200). The maximum injection time (max IT) was 200 ms at resolution of 70 000 and 450 ms at resolution of 140 000. Automated gain control (AGC) was targeted at 3 × 106 ions. For targeted MS2 analysis, the isolation width of the precursor ion was set at 1.0 (*m/z*), high energy collision dissociation (HCD) was 35%, and max IT is 100 ms. The resolution and AGC were 35 000 and 200 000, respectively.

### Metabolite Peak Extraction and Data Analysis

Raw peak data was processed on Sieve 2.0 software (Thermo Scientific) with peak alignment and detection performed according to the manufacturer’s protocol. The method “peak alignment and frame extraction” was applied for targeted metabolite analysis. An input file of theoretical *m/z* and detected retention time was used for targeted metabolite analysis, and the *m/z* width was set to 5 ppm. An output file was obtained after data processing that included detected *m/z* and relative intensity in the different samples.

## Metabolic flux analysis

### Flux model and constraints

The flux model includes biochemical reactions and diffusions between tissue and plasma. The model covers glucose, lactate and pyruvate in two or three kinds of tissue and plasma (figure 1B). All metabolites were assumed to be balanced during simulations, which means sum of income fluxes to a certain metabolite equals to sum of outgo fluxes from this metabolite. Circulatory flux constraints were implemented to reduce degrees of freedom (Hui et al., 2017).

### MID prediction, flux fitting and solution sampling

An iterative optimization algorithm was used to compute the fluxes. Given a set of flux values, MID of target metabolites can be predicted from averaging MID of precursors using flux values as weights:

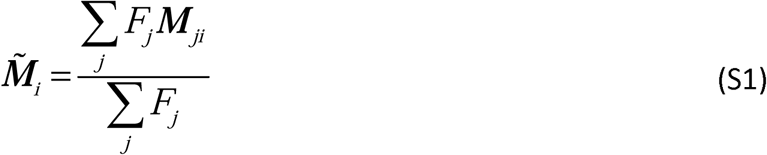

where 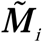 is the predicted MID of metabolite *i*, ***M*** _*ji*_ is MID of metabolite *i* produced from its precursor metabolite *j*, and *F*_*j*_ is the flux from *j* to *i*.

The difference between the predicted and experimental MIDs was evaluated by Kullback–Leibler divergence:

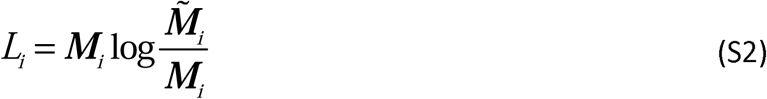

where 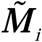 is predicted MID and ***M***_***i***_ is experimental MID of metabolite *i*. Sum of *L*_***i***_ for all target metabolites was regarded as the cost function to minimize. The optimization problem was defined as:

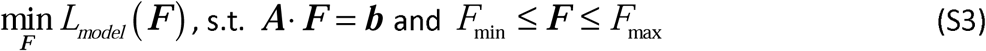

where ***F*** is the vector of all fluxes and ***A***· ***F*** = ***b*** represents flux balance requirement and other constraints.

To better cover all possible results, the high-dimensional solution space was sampled and all solutions with cost function value lower than a threshold and TCA flux value higher than a minimal requirement were selected for further calculations.

Solutions to the random unfitted control are generated using flux balance requirement and other flux constraints such as non-negative values, but without MID information.

### Distribution of flux and cost function

Values of all variable fluxes and the cost function in all feasible solutions are collected. Their distributions are displayed as a box plot, in which boxes represent median and two quantiles, and whiskers represent two extremes. Distributions of cost function are compared with that of unfitted solutions, and the pairwise p-value between cost functions of fitted and unfitted solutions is calculated based on nonparametric Wilcoxon rank-sum test.

### Glucose contribution calculation

After fitting a flux from the MID data, local and global glucose contributions were calculated to quantify the net contribution from glucose in plasma to TCA cycle. First, the contribution flux from each circulating metabolite was calculated. The contribution flux was calculated by the net flux from the plasma to each kind of tissue, minus the part which flows out of the tissue in the form of other metabolites (supplementary information). Then, the contribution flux was normalized to total contribution flux from all carbon sources to calculate the relative contribution ratio, in which the local one was normalized by contribution fluxes in sink tissue, while the global one was normalized by contribution fluxes in the complete model. All selected flux solutions from sampling process were analyzed for glucose contribution, and their relative ratios were displayed on the violin plot.

### Parameter sensitivity analysis

Experimental MID data or flux constraints were perturbed by multiplying Gaussian-distributed noises. After each perturbation, the perturbed value was used to solve fluxes and calculate the local distribution of glucose contribution. Then the median value of the distribution was calculated. Median values from all perturbations were collected and displayed by violin plot.

### Hypoxia correction analysis

Experimental MIDs of glucose in source tissue was assumed to be a mixture of 80% original glucose MID and 20% hydrolyzed glucose from glycogen (unlabeled MID), and the MID of lactate in the sink tissue was assumed to be a mixture of the original lactate MID and 20% a product from pyruvate (same MID as pyruvate in sink tissue). Therefore, the putative original MID can be solved and used for the same fitting and calculation of glucose contribution ratio.

### Ternary graph plotting

Contributions from three energy sources were visualized by ternary graph. All triplets of contribution ratios were plotted on the triangle and density of distribution was rendered by a Gaussian kernel. In the rendering process, convolutions between all points and Gaussian distribution were calculated, and the resulting density was converted into space of ternary graph to be displayed.

### Software implementation

Scripts in this study were implemented by Python 3.6. Source codes are available from GitHub (https://github.com/LocasaleLab/Lactate_MFA). The package version dependency is also provided on GitHub website. A Docker on Linux system for out-of-the-box running is also available. Each model requires around 10 ∼ 50 hours of running time.

## ACKNOWLEDGEMENTS

We thank members of the Locasale laboratory for helpful discussions. Support from National Institutes of Health (R01CA193256, R35CA197616) and the American Cancer Society (RSG-16-214-01-TBE) are gratefully acknowledged. JWL has advisory roles for Nanocare Technologies, Raphael Pharmaceuticals, and Restoration Foodworks.

## AUTHOR CONTRIBUTIONS

S.L., Z.D., and J.W.L. designed the study. S.L. and J.W.L. wrote the manuscript with essential input from Z.D. S.L. developed the model and performed the data analysis with help from Z.D. D.E.C. performed *in vivo* nutrient infusions and reviewed results with D.G.K. All authors have read, edited and approved the final manuscript.

## LEGENDS

**Figure S1.**
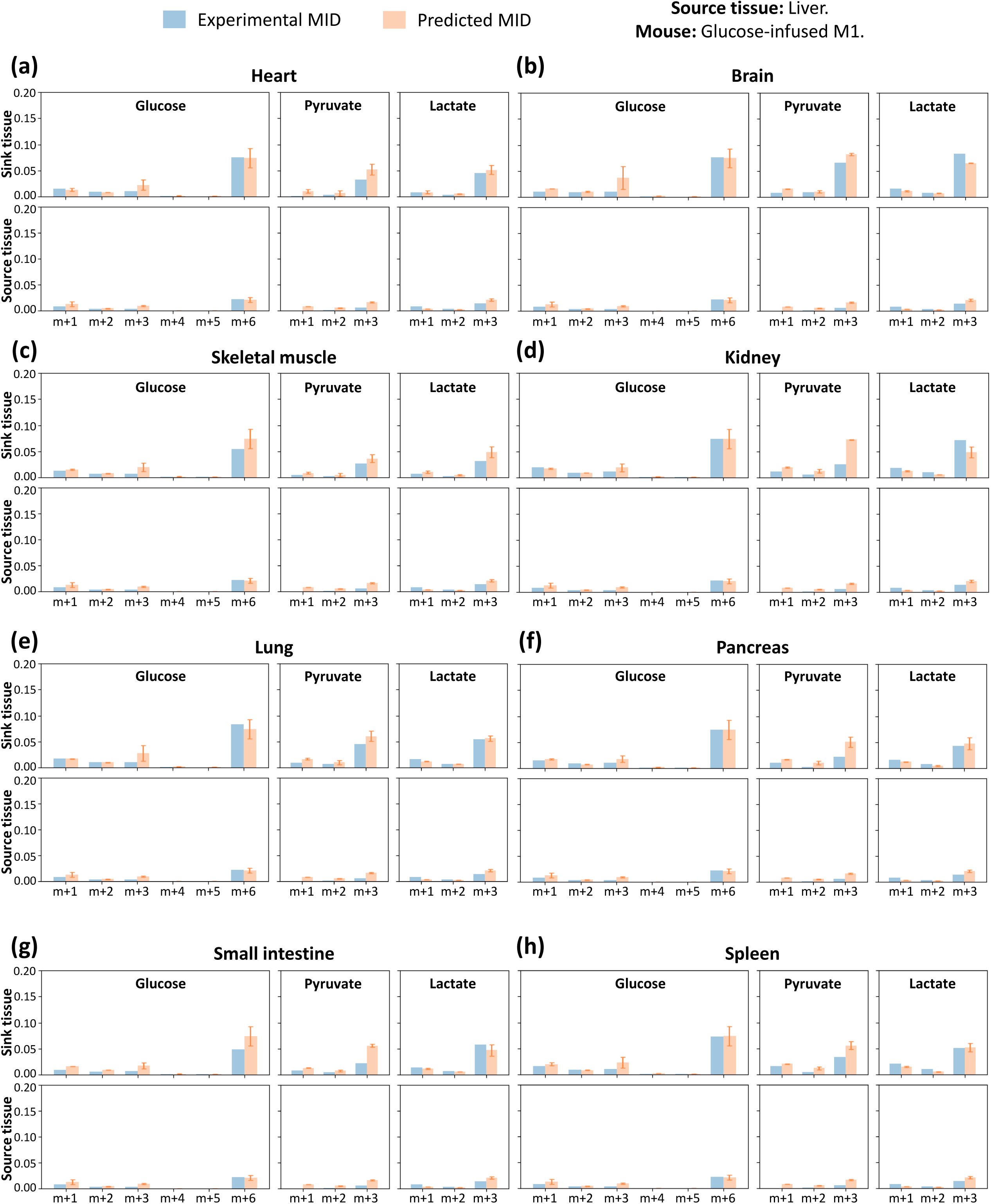
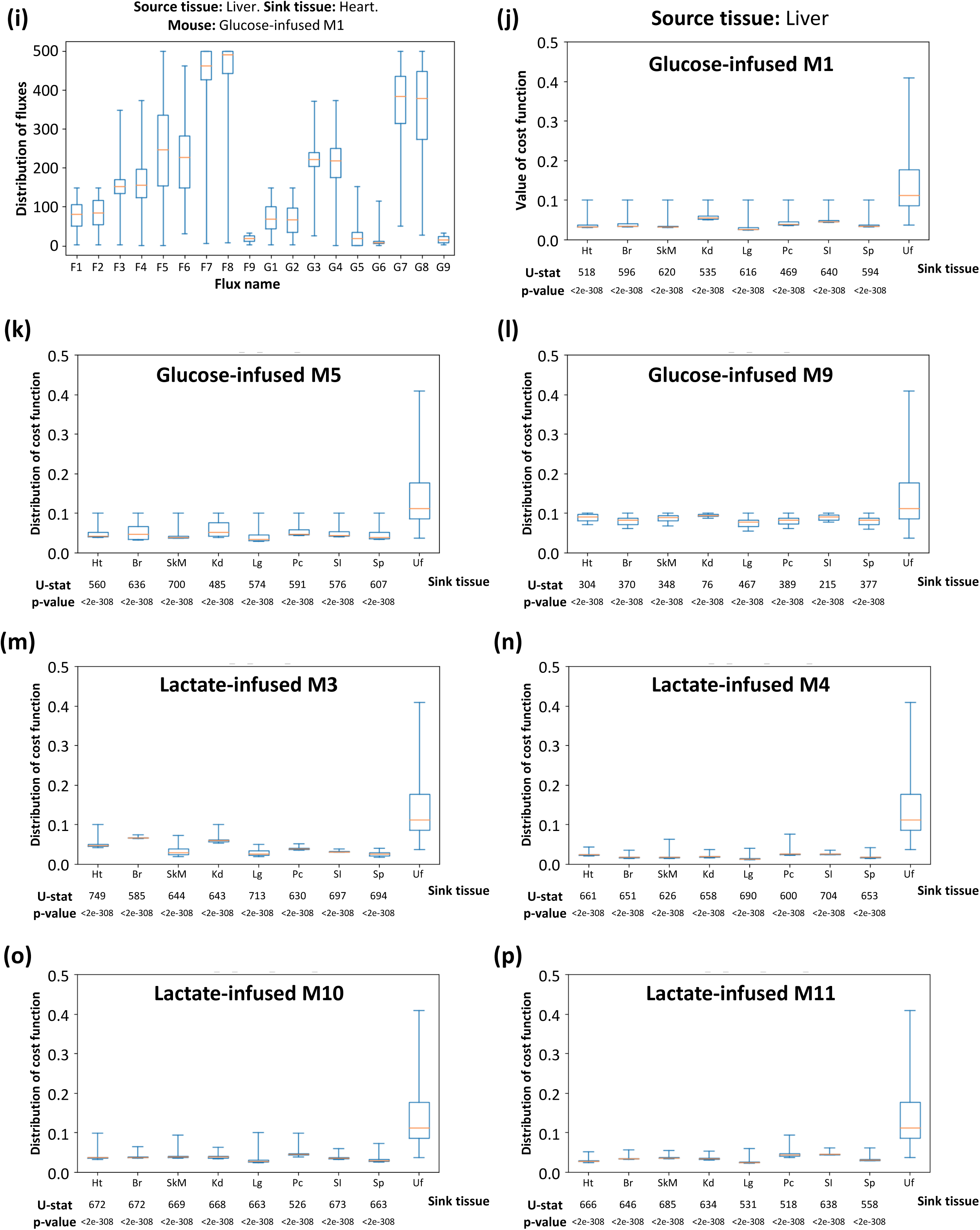
Detailed information of model fitting. (a-h) Comparison of experimental and predicted MID in the model. Average values of predicted MID in all feasible solutions are displayed. Standard deviation is also displayed as error bar. In most cases, the experimental MID is captured by this model. Because of the low abundance of labeled isotopomers, only isotopomers with more than one ^13^C are displayed. (i) Distribution of 18 variable fluxes in all feasible solutions. Source tissue is liver and sink tissue is heart. (j-p) Distribution of cost functions of feasible solutions fitted with different sink tissue and data from different mice, compared with unfitted control data. Source tissue in all fittings is liver. U-statistics of rank-sum test and p-values are displayed. In all subfigures, fits are based on mice from the low-infusion data in Hui et al, 2017, and source tissue is liver. Specifically, subfigures (a-h) and (i) are fitted with glucose-infused mouse M1. The sink tissue in (i) is heart. In all box plots, boxes represent quantiles and whiskers represent extremes. Ht: heart, Br: brain, SkM: skeletal muscle, Kd: kidney, Lg: lung, Pc: pancreas, SI: small intestine, Sp: spleen, Uf: data from random unfitted control.

**Figure S2.**
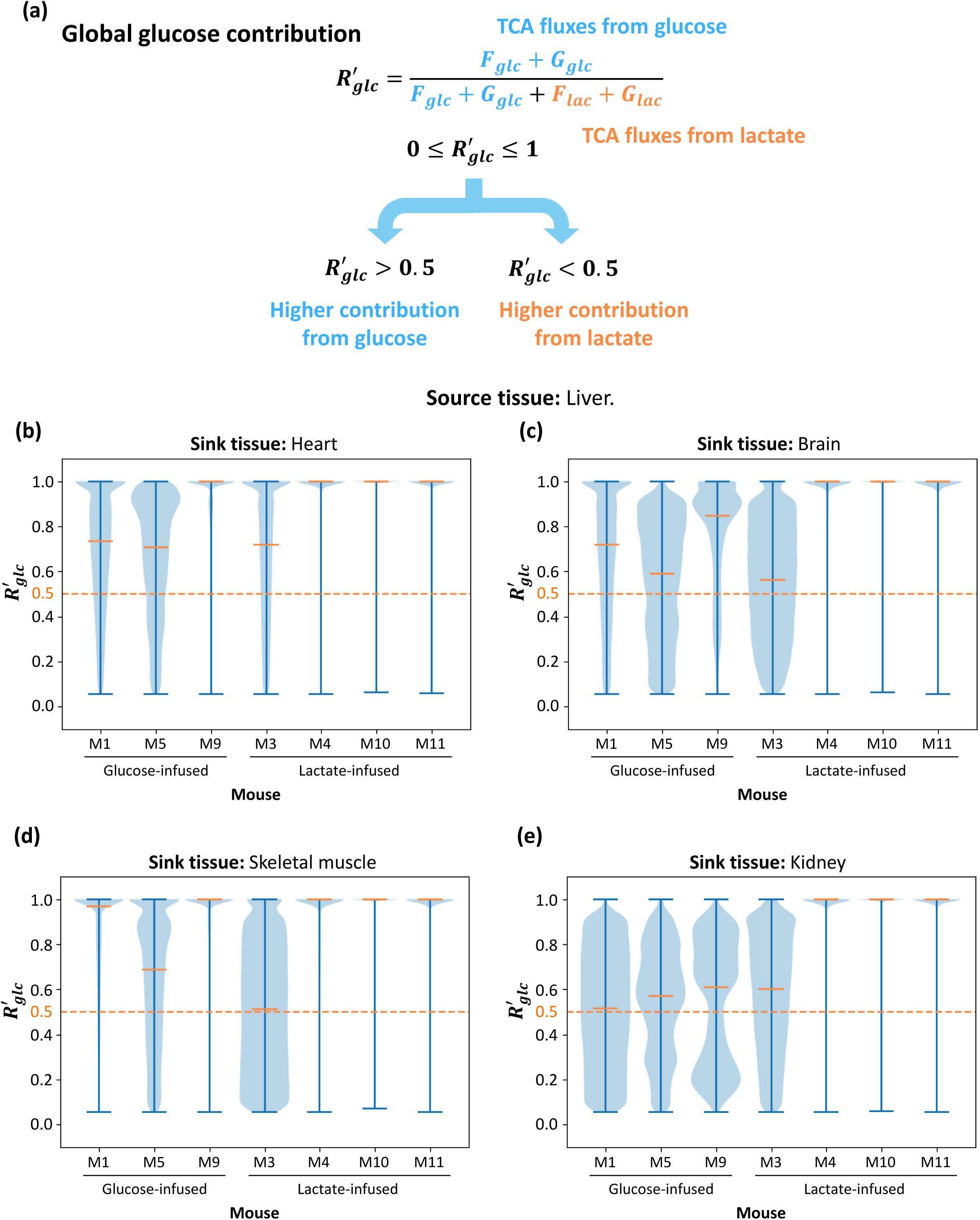
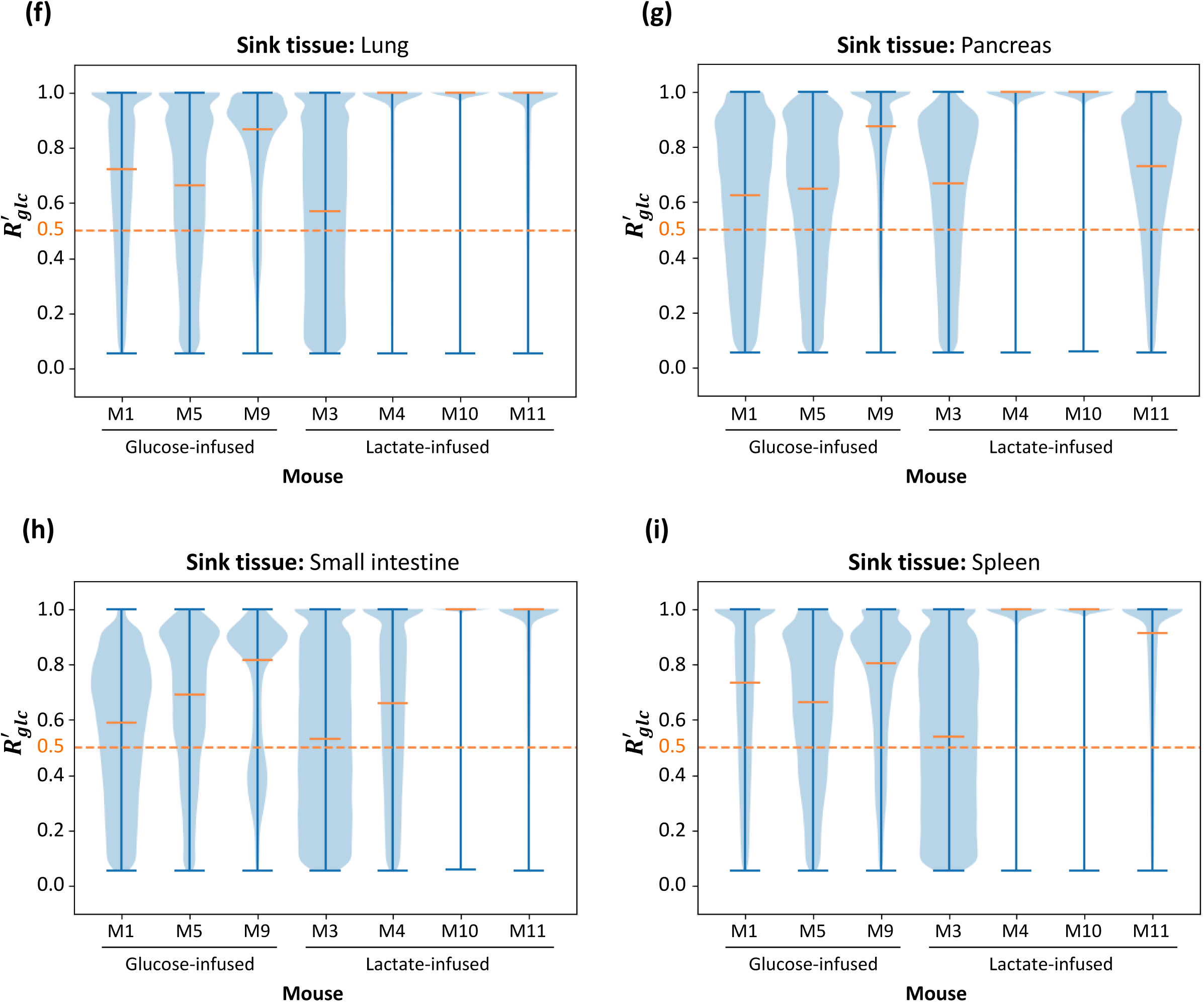
The global contribution ratio from circulating glucose to the TCA cycle. (a) Definition of global glucose contribution ratio in complete model 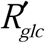. The global contribution 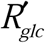 is defined as the relative ratio of glucose contribution flux to total contribution flux in sink and source tissue. Similar with *R*_*glc*_, this contribution is also a scalar between 0 and 1. Higher 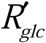 represents an increasing glucose contribution to the TCA cycle. (b-i) Distribution of glucose contribution in complete model based on the model with different sink tissues. For each sink tissue, the source tissue is liver and contribution ratio is calculated from data in 7 different mice. Similar with results with *R*_*glc*_, for most kinds of sink tissue, the median of global glucose contribution 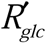 is close to or higher than 0.5 in most mice, which means glucose contributes more than lactate to the TCA cycle in complete model. The orange dash line represents 0.5 threshold. Data set is from glucose-infused mice (M1, M5, M9) and lactate-infused mice (M3, M4, M10, M11) in Hui et al, Nature 2017.

**Figure S3.**
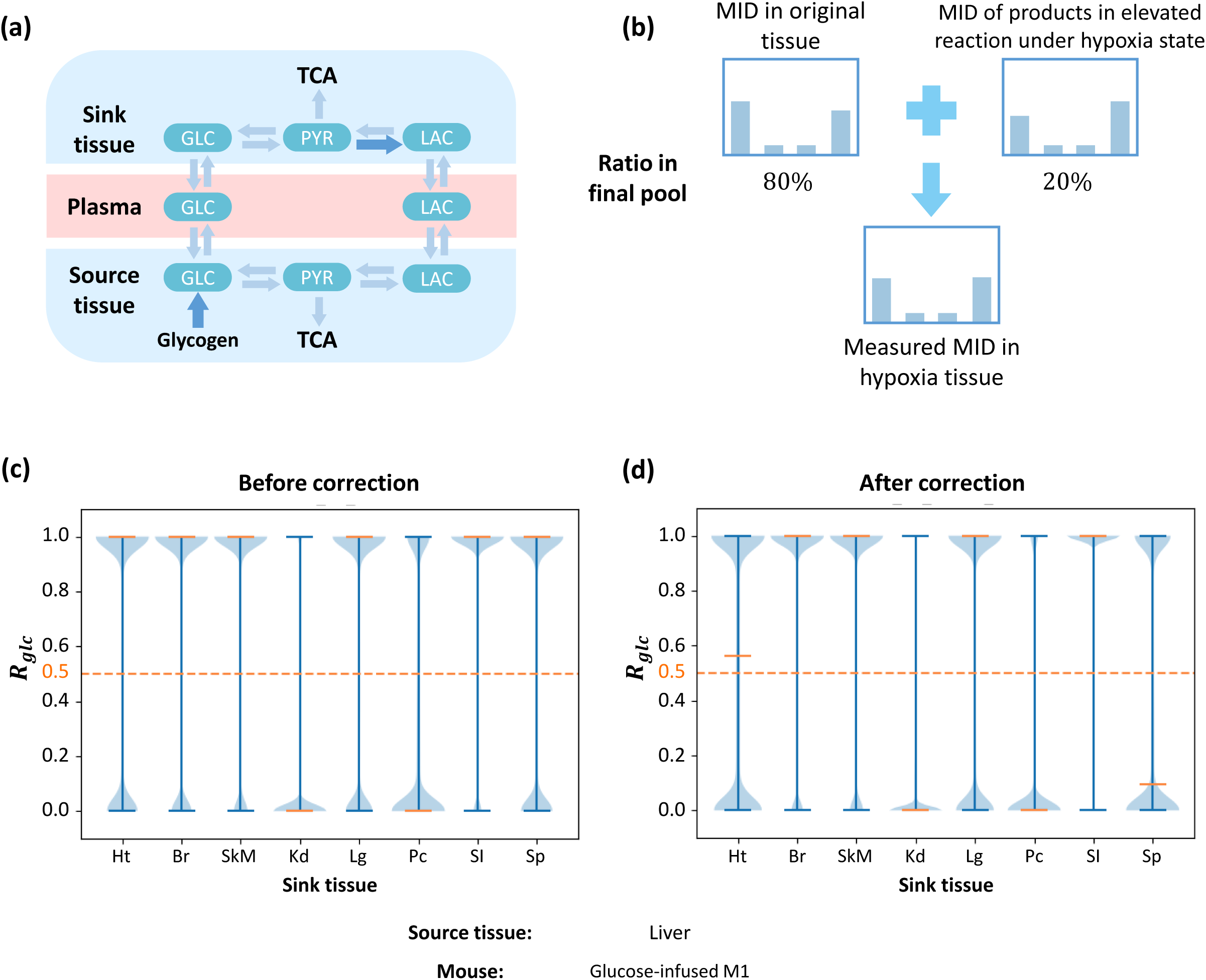
Distribution of local glucose contribution under hypoxia correction. (a) Two possible elevated fluxes under hypoxia. (b) Final measured MID is regarded as mixture of MIDs from real metabolite and products in elevated reaction under hypoxia state. (c-d) Distribution of local glucose contribution before (c) or after (d) correction. Model is fitted with data from mouse M1 in data from Hui et al Nature 2017. Sink tissue is liver. Ht: heart, Br: brain, SkM: skeletal muscle, Kd: kidney, Lg: lung, Pc: pancreas, SI: small intestine, Sp: spleen.

**Figure S4.**
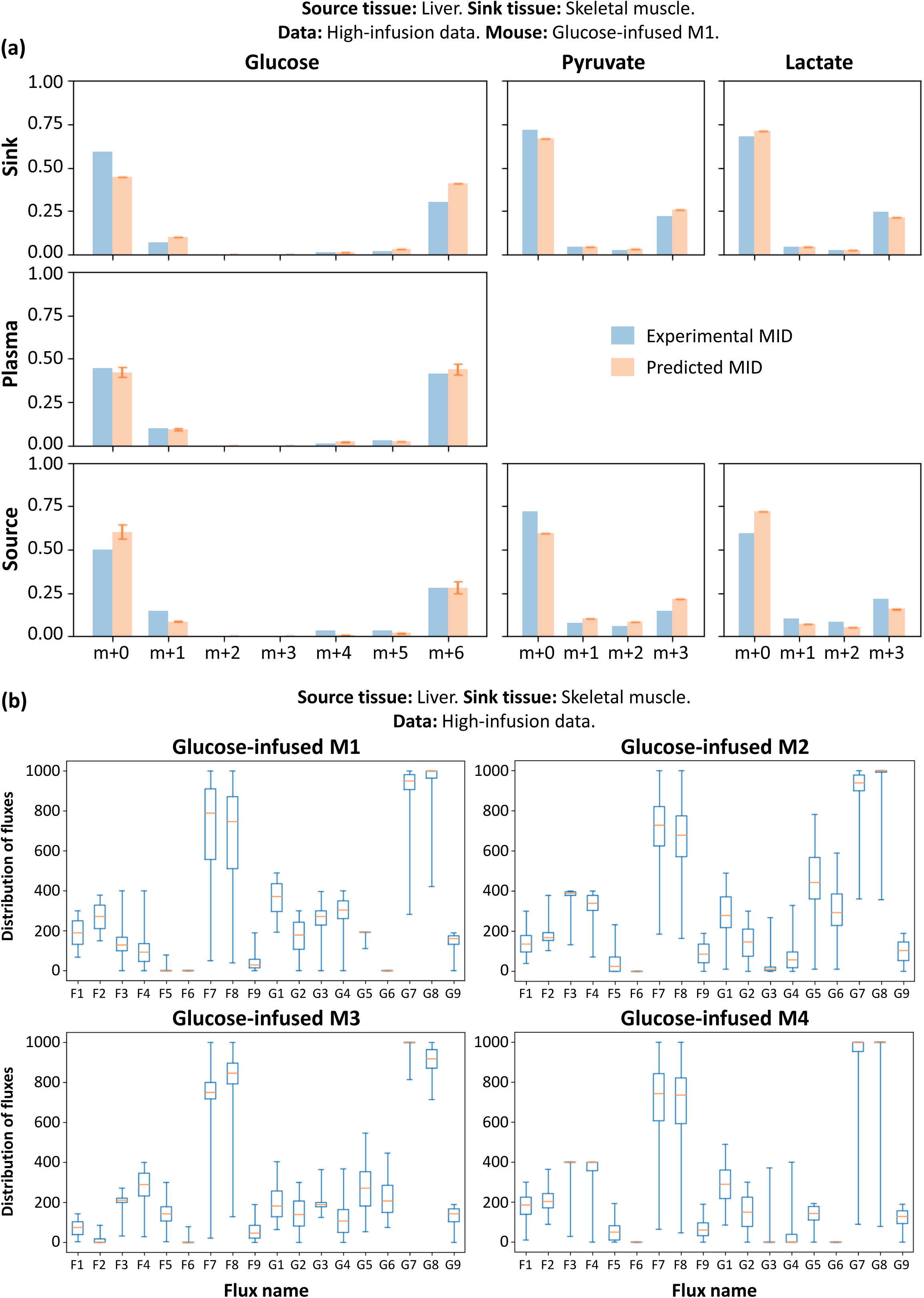
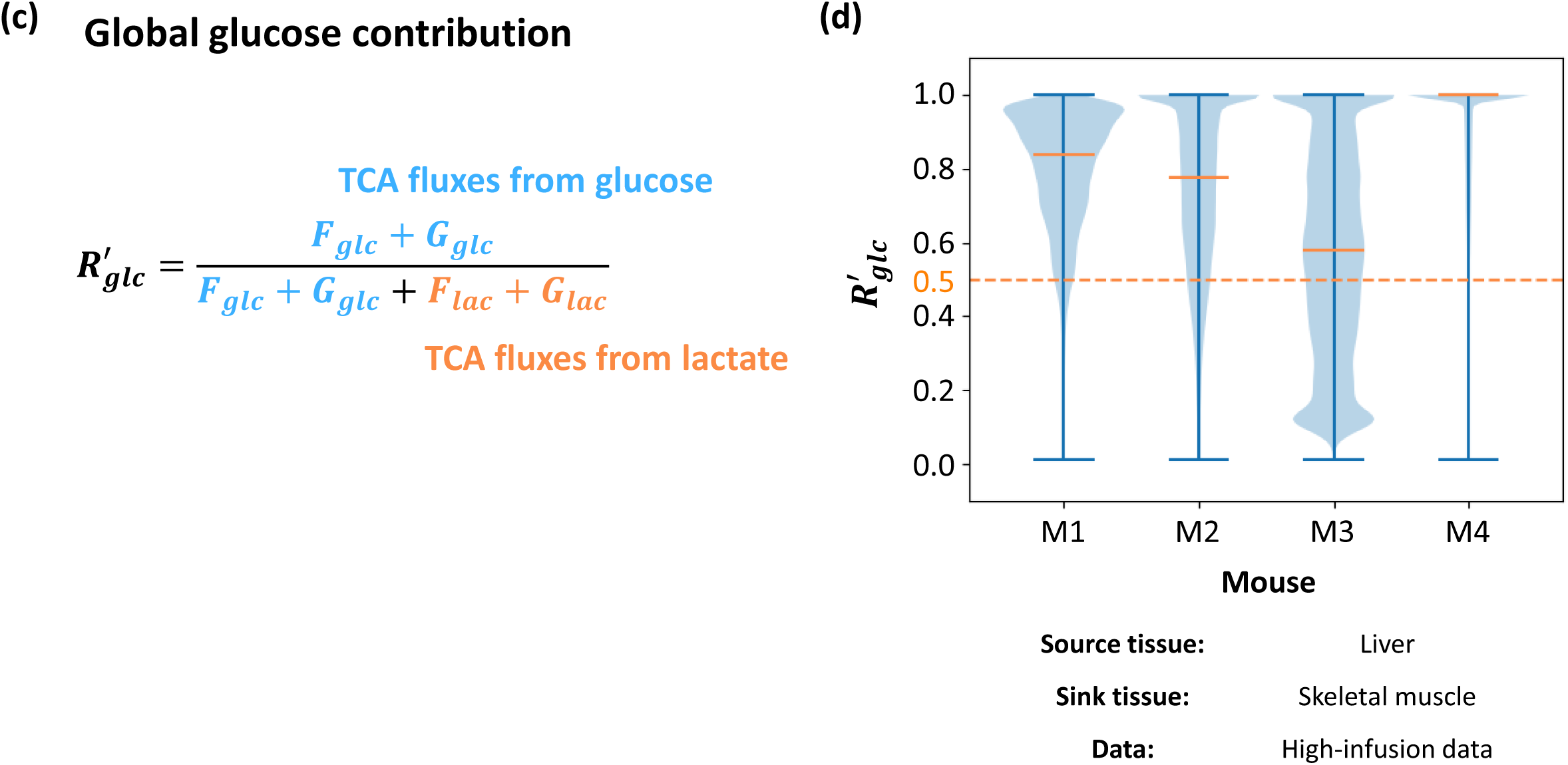
Detailed information for the high-infusion system. (a) Comparison of experimental and predicted MID in the model. Average of predicted MID in all feasible solutions are displayed. Standard deviation is also displayed as error bar. Experimental MID could be well predicted by this model. Notably, the ratio of high ^13^C isotopomer is significantly higher than that of the original system. (b) Distribution of 18 variable fluxes in all feasible solutions. (c) Definition of global glucose contribution in complete model 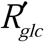. The global glucose contribution 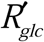 is defined as the relative ratio of glucose contribution flux to total contribution flux in sink and source tissue. (d) Similar with results with local contribution *R*_*glc*_, distribution of global glucose contribution in complete model 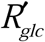 shows glucose contributes more than lactate to the TCA cycle in all mice. Fittings from different mice are displayed. In all subfigures, the fitting is based on different glucose-infused mice from the high-infusion data. Source tissue is liver and sink tissue is skeletal muscle. Specifically, subfigure (a) is based on data from glucose-infused mouse M1. In all box plots, boxes represent quantiles and whiskers represent extremes.

**Figure S5.**
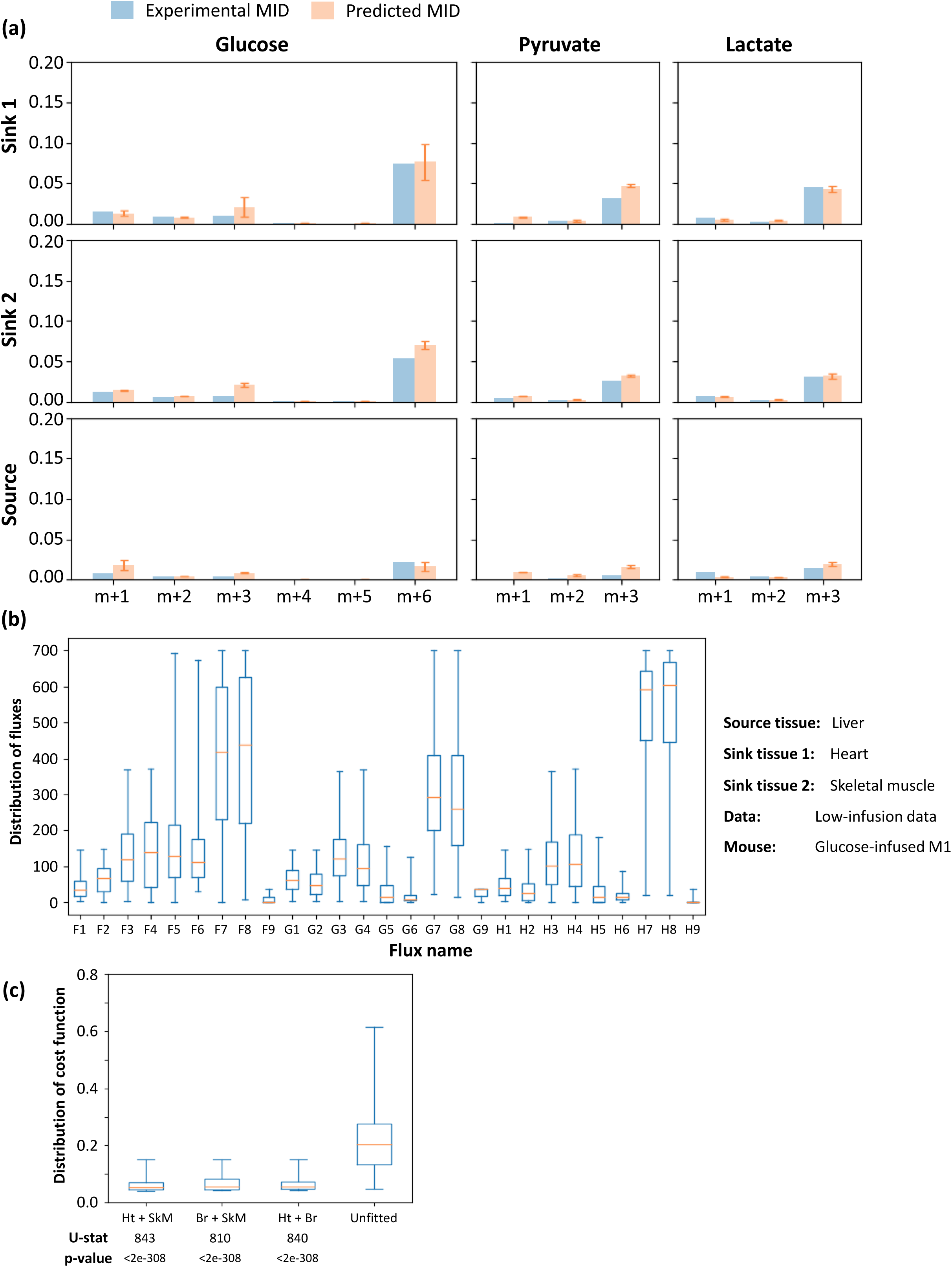
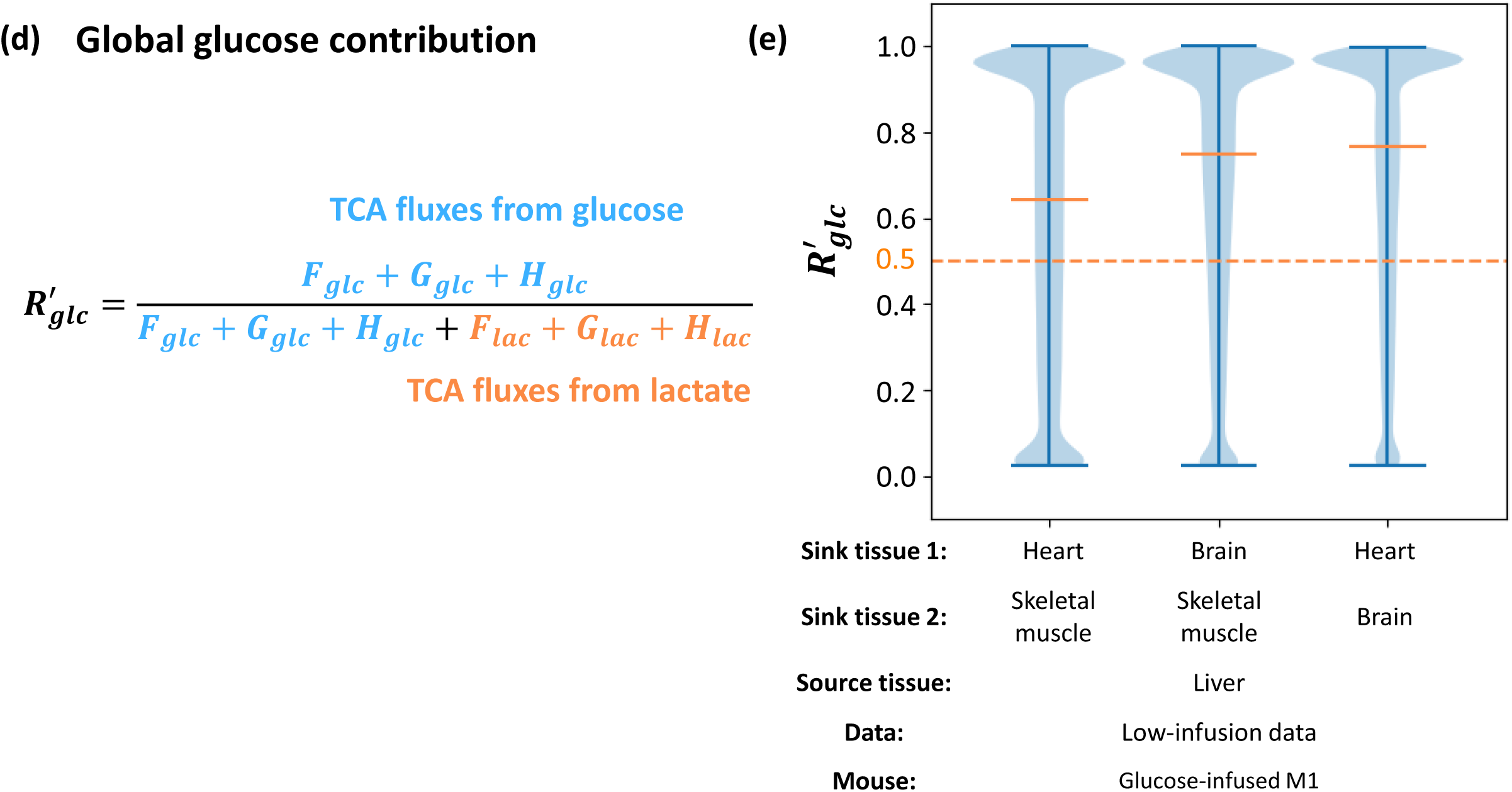
Detailed information of the multi-tissue model. (a) Comparison of experimental and predicted MID in the model. Average of predicted MID in all feasible solutions are displayed. Standard deviation is also displayed as error bar. Experimental MID could be well predicted by this model. Because of the low abundance, only isotopomers with more than one ^13^C are displayed. (b) Distribution of 27 variable fluxes in all feasible solutions. (c) Distribution of cost functions of feasible solutions compared with that from unfitted control. U-statistics of rank-sum test and p-values are displayed. (d) Definition of global glucose contribution in complete model 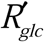. This global contribution 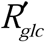 is defined as the relative ratio of glucose contribution flux to total contribution flux in all three kinds of tissue. (e) Similar with results with local contribution *R*_*glc*_, distribution of global glucose contribution in complete model 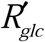 shows glucose contributes more than lactate to the TCA cycle in all combinations of sink tissue. In all subfigures, the model is fitted with glucose-infused mouse M1 from the low-infusion data in Hui et al, 2017. The source tissue is liver and the sink tissue 1 and 2 are two from heart, brain and skeletal muscle respectively. In all box plots, boxes represent quantiles and whiskers represent extremes.

**Figure S6.**
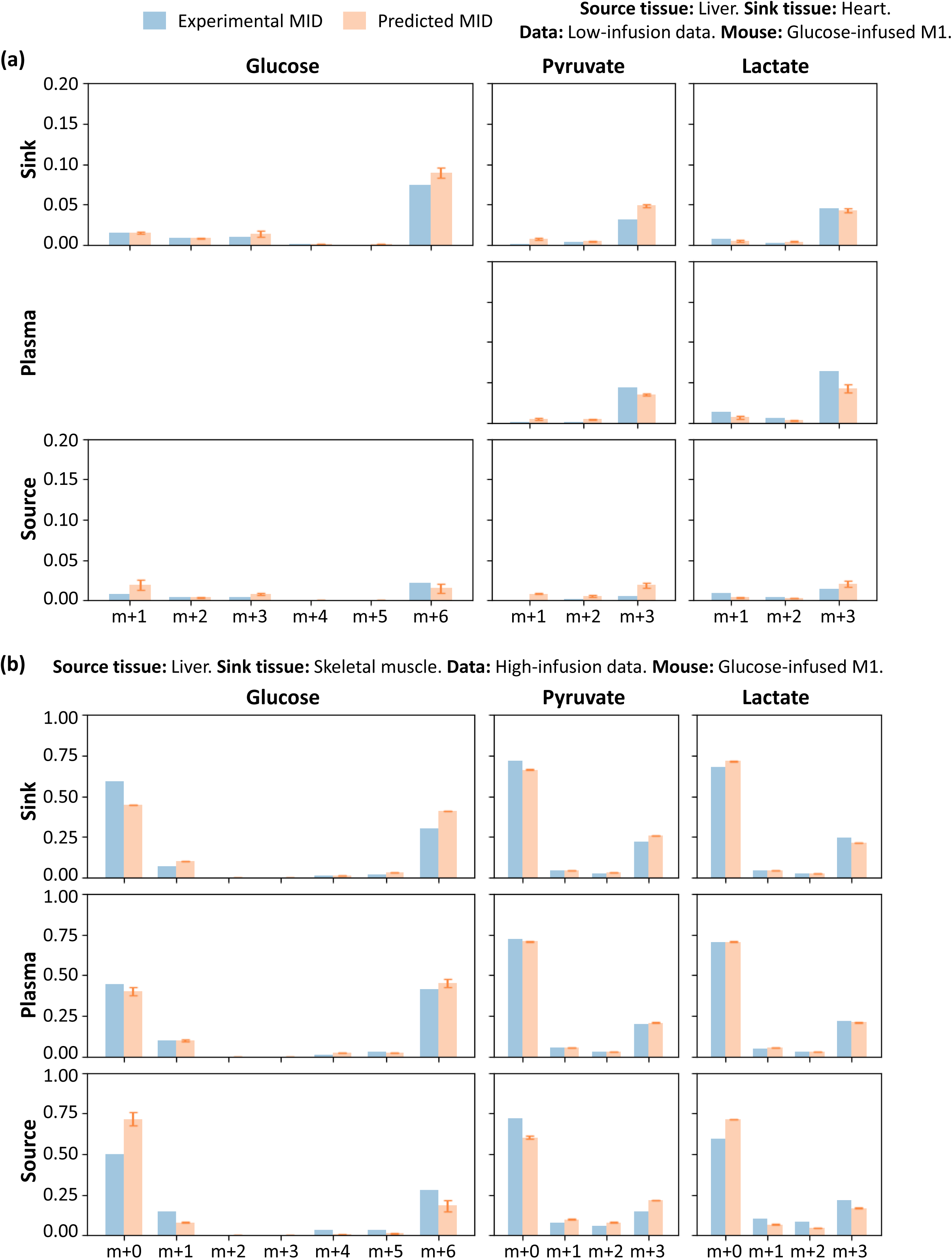
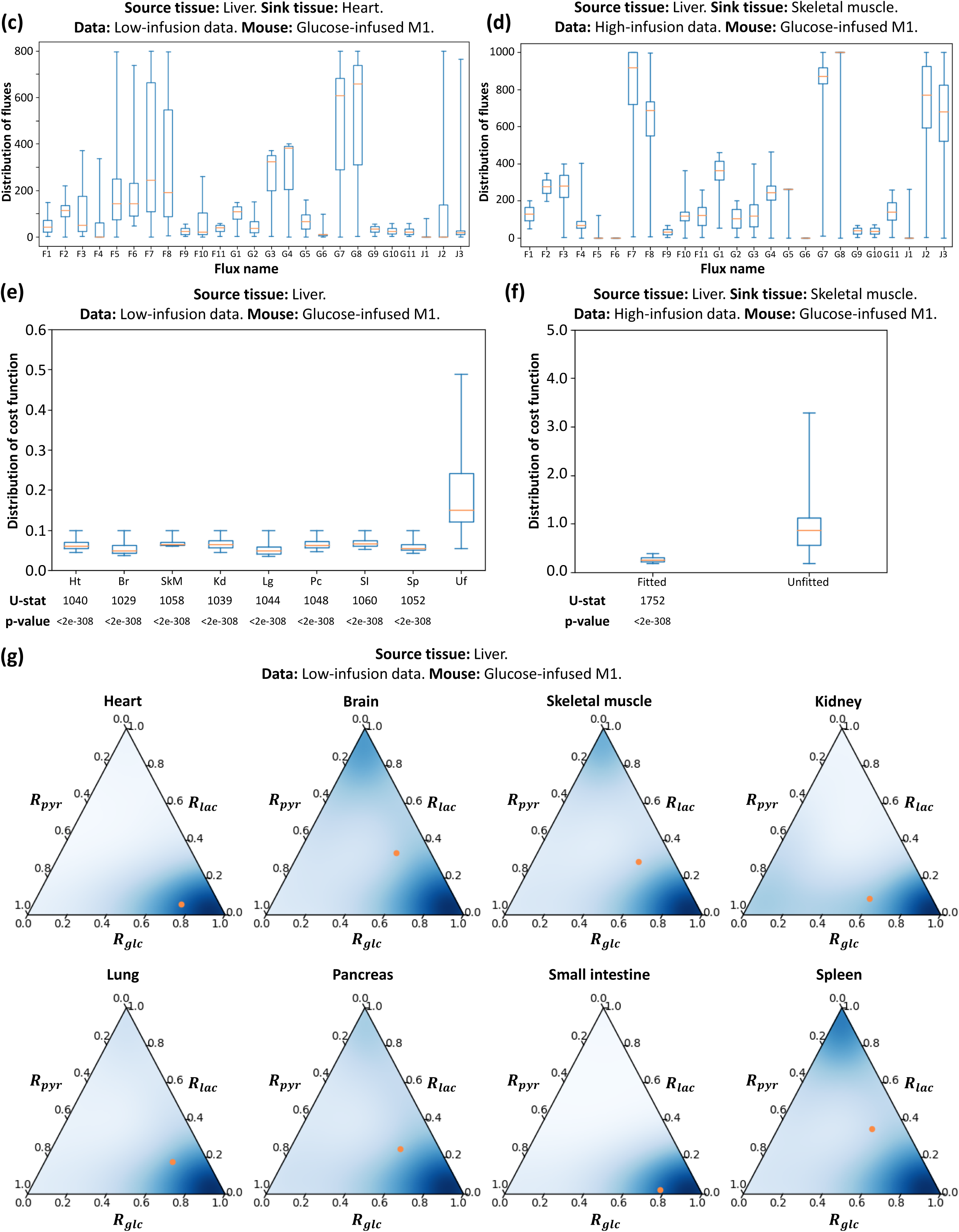
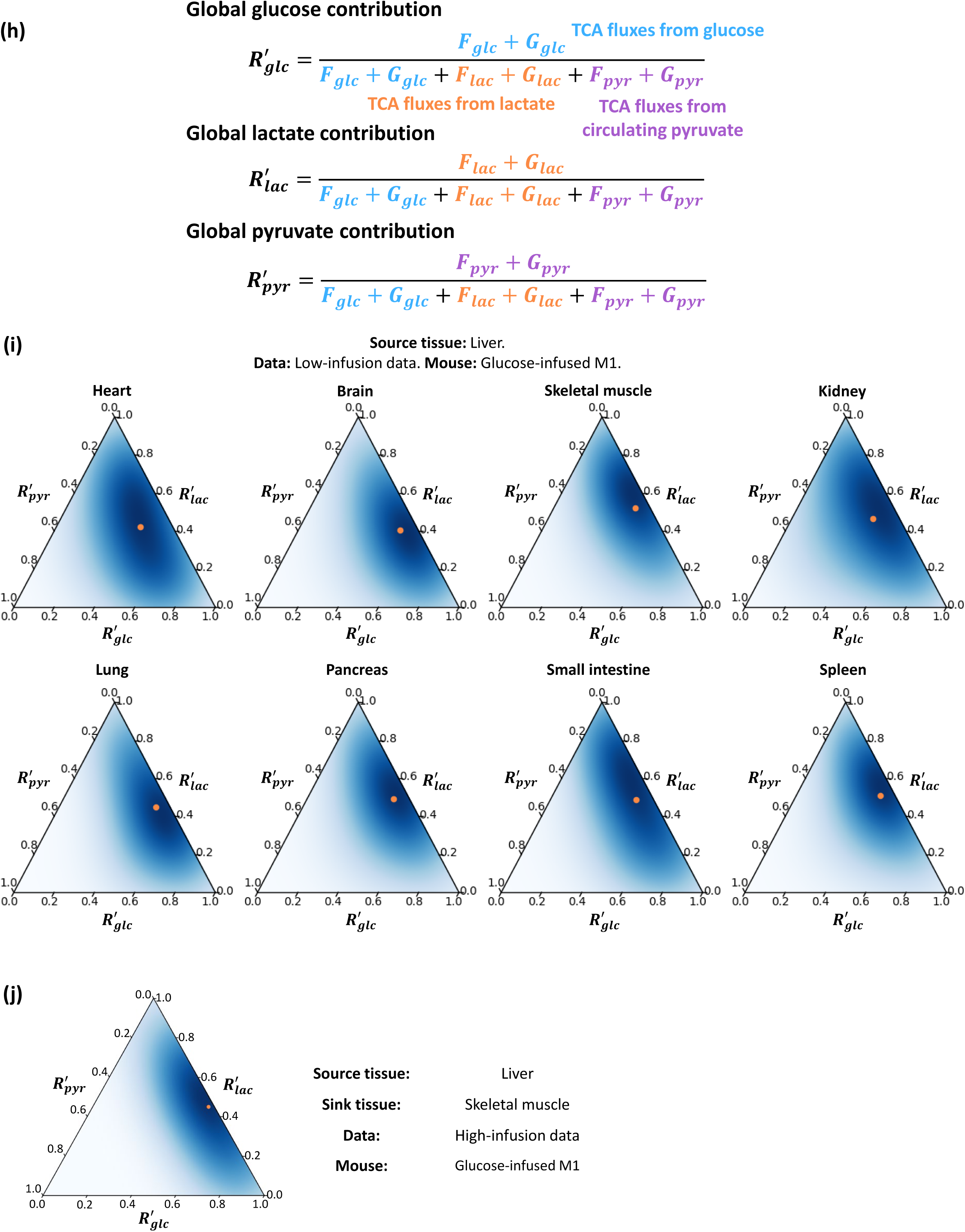
Detailed information of the model with multiple circulating metabolites. (a, b) Comparison of experimental and predicted MID in the model fitted with the low-infusion data (a) or the high-infusion data (b). Average of predicted MID in all feasible solutions are displayed. Standard deviation is also displayed as error bar. Experimental MID could be well predicted by this model. (c, d) Distribution of 25 variable fluxes in all feasible solutions fitted with the low-infusion data (c) or the high-infusion data (d). (e, f) Distribution of cost functions of feasible solutions compared with that from random unfitted control, fitted with the low-infusion data (e) or the high-infusion data (f). U-statistics of rank-sum test and p-values are displayed. (g) Local contributions to the TCA cycle when additional nutrients are considered. For most sink tissues, glucose contributes to the TCA cycle more than lactate and pyruvate, especially in heart, kidney, lung, and small intestine. The orange point indicates average level. (h) Definition of global contribution from metabolites in complete model 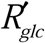, 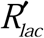 and 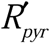. The three global contribution ratios 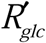, 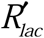 and 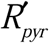 are defined by the relative ratio of the contribution flux from each metabolite to total contribution flux of all three metabolites in sink and source tissue. (i) Ternary plot to display distribution of three contributions 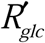, 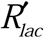 and 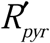. The orange point indicates average level. (j) Analysis and results as in (i) but for additional high-infusion system of different animal strain, different diet and different infusion protocol. Subfigure (a), (c), (e), (g) and (i) are fitted with glucose-infused mouse M1 from the low-infusion data in Hui et al, Nature 2017, and source tissue is liver. Specifically, sink tissue in (a) and (c) is heart. Subfigure (b), (d), (f) and (j) are fitted with glucose-infused mouse M1 from the high-infusion data, and the sink tissue and source tissue are liver and skeletal muscle respectively.

