## Supplementary Methods for "Quantitative analysis of the physiological contributions of glucose to the TCA cycle"

**Supplementary Information for “Quantitative analysis of the physiological contributions of glucose to the TCA cycle”**

### General Method

#### Model design

A principle of model design is parsimony, also referred to as “Occam’s razor” that is to use the simplest model that can appropriately address the question at hand. Inclusion of parameters and variables should correspond to available data and constraints that parameterize the model and allow for the model to address the relevant questions. The primary goal of this study is to quantify the contribution of circulating glucose and lactate to the TCA cycle. Therefore, the model, to reach appropriate conclusions, should balance the ability to achieve this goal and the complexity that can be precisely evaluated from available data.

To study circulating metabolites, the model contains at least two compartments: plasma and a specific tissue. However, because circulating glucose and lactate must be balanced, the net flux between plasma and tissue is limited to boundary fluxes of circulating glucose and lactate, which cannot capture dynamics between circulation and target organs. Therefore, a heterogenous two tissue system is introduced, and it allows for different patterns in utilization of nutrient source. One common pattern is the Cori cycle, in which in the fasting state, the sink tissue (muscle) utilizes circulating glucose and excretes lactate, while the source tissue (liver) convert them back to glucose.

The basic structure using two tissues and plasma is already difficult to model. To prevent overfitting leading to parameter uncertainty, the network in each tissue contains three key nodes: glucose, pyruvate and lactate, and their interconversion fluxes. TCA-related reactions are described by one unidirectional flux, because introducing more TCA cycle reactions would not address the question of relative nutrient contributions and lead to overfitting. To intuitively understand whether TCA-related reactions have a substantial impact on the labeling patterns, the MID of phosphoenolpyruvate (PEP), the metabolite generated from oxaloacetate (OAA) in the first step of gluconeogenesis, is also measured in liver (source tissue) and skeleton muscle (sink tissue) in the high-infusion rate data. The MID for PEP is uncorrelated with that of malate in TCA cycle (Schematic 1b), suggesting that the cataplerotic flux from the TCA cycle intermediates to glucose (which requires PEP as an intermediate) does not have a significant impact on labeling patterns of metabolites. Similar results are also observed in a previous study from Hui et al Nature 2017. Although the MID of PEP is not available in those data, 3-phosphoglycerate (3PG), the metabolite adjacent to PEP in glycolysis/gluconeogenesis (Schematic 1a), showed a similar trend, again indicating that the effect of the labeling pattern from TCA cycle intermediates to glucose is very low. Therefore, although previous studies show that cataplerotic flux of TCA may be one of major sources of PEP in liver, introducing more TCA reactions do change results of the fits from current metabolites, and thus not significantly improve the precision of this model.

However, a more complicated model that includes every reaction carrying the fluxes of the TCA cycle, at least 6 metabolite MIDs and tens of fluxes should be added, including citrate, α-ketoglutarate (as well as glutamate), succinate, oxaloacetate (as well as aspartate) in the TCA cycle and PEP in glycolysis. Absolute measurements of fluxes feeding into the TCA cycle from other carbon sources such as branched chain amino acids and glutamine are also required to fully parameterize the model. The lack of data would lead to overfitting and parameter uncertainty which limits the conclusions that can be drawn. Under this condition, introducing more detailed fluxes may not substantially improve fitting precision, but would introduce uncertainties within the current model. Therefore, after careful consideration of the available data and the primary goal of the model, the resulting model consists of one plasma and two tissues, which includes glucose, pyruvate, lactate and conversion fluxes between them.

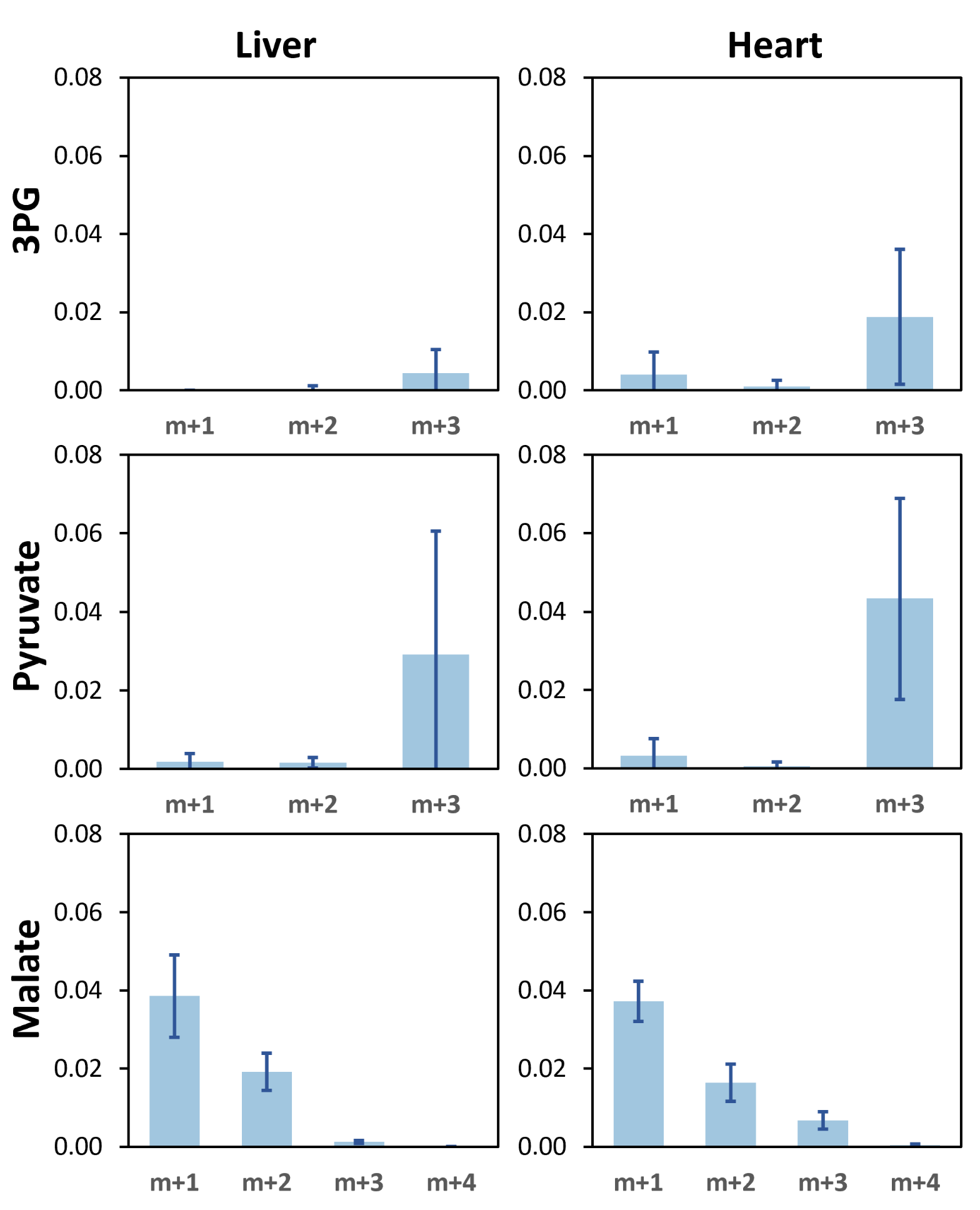

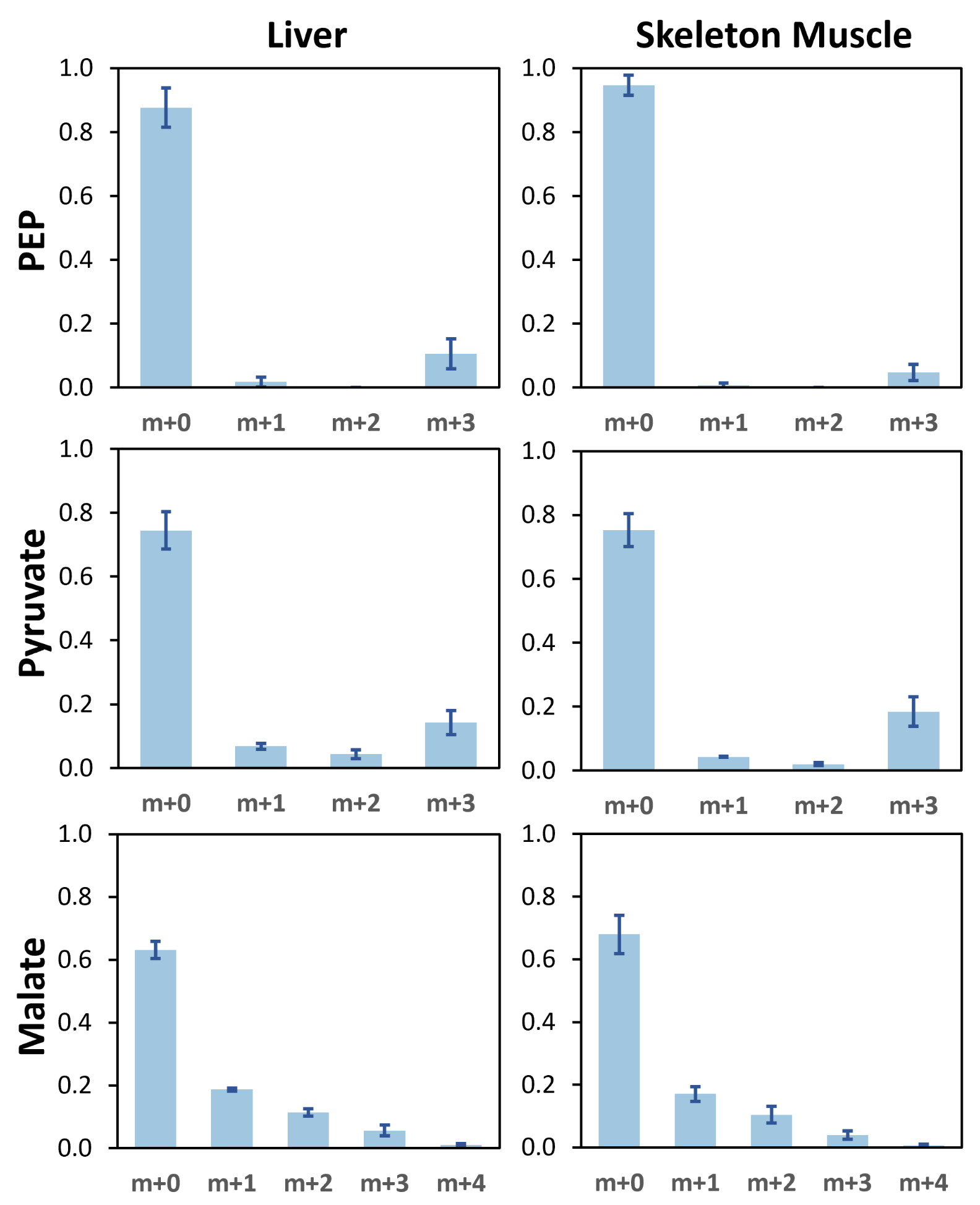

**(a)**

**(b)**

**Schematic 1**. MID data of metabolites. Error bars indicate standard deviations. (a) MID of 3PG, pyruvate and malate in liver and heart in glucose-infused data from Hui et al Nature 2017. Data for 3PG and malate were collected from 5 mice (labeled as M1, M4, M5, M7 and M9 in their experiments), while those for pyruvate were collected from 3 mice (M1, M5 and M9. Data for pyruvate M4 and M7 were not available). Only non-m+0 isotopomers are plotted because of low abundance. (b) MID of PEP, pyruvate and malate in liver and skeleton muscle in glucose-infused data from our high-infusion experiments. Data were collected from 4 mice.

There are some limitations in the current model. For example, in the source tissue, M+3 PEP is higher and M+3 pyruvate is lower compared with that in sink tissue. considering the similar abundance of M+6 glucose both in source and sink tissue (Figure S4A), this implies that there might be more high-labeled carbon source that supply of PEP in the source tissue. Limited by available data, the current model cannot explain the source of those hidden high-labeled carbon sources. Similarly, lower M+3 pyruvate in the source tissue might be due to some unlabeled sources of pyruvate in source tissue, such as glucogenic amino acids. However, data also show that the abundance of M+4 in malate in liver is very low, both in the Hui et al. Nature 2017 data and the high-infusion data. Considering the high exchange rate between malate and aspartate/oxalacetate, although the cataplerotic flux may be one of the main sources of PEP, it should not be the main reason for higher M+3 PEP in source tissue, nor the main reason of the difference between experimental and predicted MID in this study.

The structure of this model is another point of discussion. For example, in cellular metabolism, the flux G2 relies on glucose 6-phosphatase (G6Pase), but most kinds of sink tissue lack this enzyme. Similarly, the flux G6 relies on phosphoenolpyruvate carboxykinase (PEPCK), which is often thought to be most highly expressed in liver. However, these two fluxes are both preserved in the sink tissue, since one of the main goal of this model is to introduce exchange fluxes of some metabolites between tissue and circulation. Exchange fluxes may significantly affect MID data, which will also influences the results of flux analysis. In tissue level, these exchange fluxes are derived as an abstracted model of a series of complicated biochemical reactions, including transport in extracellular fluid, absorption/secretion by tissue cells, and utilization/production by the tissue cells. This complicated process is not directly equivalent to a cellular biochemical reaction. On the other aspect, for the G2 flux, many sink tissues, such as kidney, heart, lung, head and leg, has been reported to have release flux or nearly release flux of glucose by direct flux measurement in pigs (Figure 3B, 3C in (Jang et al., 2019)). For the G6 flux in sink tissue, from the distribution in the region of feasible solutions, G6 is very small in most cases (Figure S1I, M1 and M4 in Figure S4B, Figure S5B, Figure S6C-D). However, because of heterogeneity in biological organisms, in some cases G6 is still very high (M2 and M3 in Figure S4B). These results indicate that the current model may reflect the difference between cellular level and tissue/organ level metabolism.

#### Flux model

In this study, each metabolic reaction network includes many fluxes between different metabolites (chemical reactions) or metabolites traversing through different tissues (diffusions). Flux models include glucose, lactate and pyruvate as metabolites in three compartments (plasma, source tissue and sink tissue). Each model contains tens of fluxes, labeled with
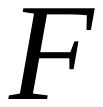
 (for fluxes in the source tissue),
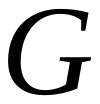
 (for fluxes in the sink tissue),
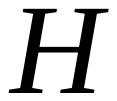
 (for fluxes in the second sink tissue in models with multiple sink tissues) and
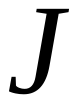
 (for fluxes within plasma). The solution is a vector
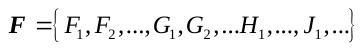
 containing flux values for all reactions in the metabolic reaction network.

It is required that all fluxes satisfy mass balance constraint, which is:

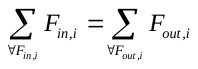
(S1)

in which
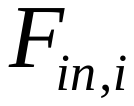
 and
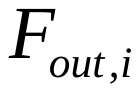
 represent all input and output fluxes connected to metabolite
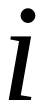
 respectively (Schematic 2). All fluxes are required to be within a range
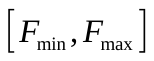
.

**Schematic 2**. Diagram of flux balance requirement. Sum of all input fluxes should be the same as sum of all output fluxes.

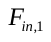

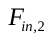

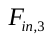

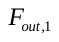

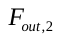

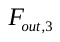

#### Flux constraints

To reduce the degrees of freedom, constraints are introduced: first, the flux that supplements glucose in the source tissue (referred as
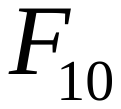
 in all models) is set as a fixed value
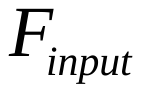
. Second, the sum of total input fluxes to plasma glucose, which is glucose turnover flux in plasma, is set as a fixed value
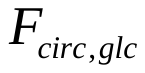
. Similarly, the lactate turnover flux in plasma is set as a fixed value
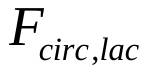
 (Schematic 3). Their values were chosen from previous research (Hui et al., 2017), and sensitivity with respect to changes in their values was evaluated.

To search the solution space, some fluxes are set to a fixed value during the fitting process. Details are explained in “Solution space sampling” section.

**Source tissue**

**Sink tissue**

**Plasma**

**TCA**

**TCA**

**Input**

GLC

PYR

LAC

GLC

GLC

PYR

LAC

**Glucose flux sum**

LAC

**Lactate flux sum**

**Input flux**

**Schematic 3**. Constraints for some fluxes are shown in orange text. “Glucose flux sum” shows the sum of two glucose export fluxes from tissues to plasma, while “lactate flux sum” shows sum of lactate fluxes. “Input flux” shows the fixed incoming glucose flux.

#### Mass isotopomer distribution calculation

The predicted mass isotopomer distribution (MID) of a metabolite is calculated based on MID of its precursors and corresponding flux values, which can be expressed as:

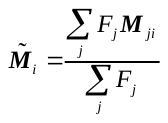
(S2)

where
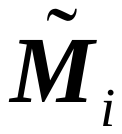
 is predicted MID of metabolite
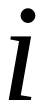
,
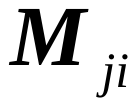
 is MID of metabolite
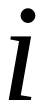
 produced from a substrate
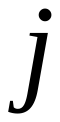
, and
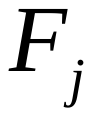
 is the flux from
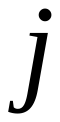
 to
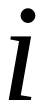
.
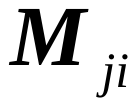
 could be calculated by experimental MID of metabolite
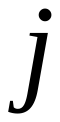
:
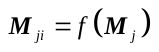
, in which
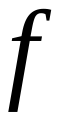
 is the MID conversion function between substrate
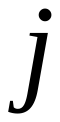
 and product
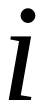
.

For example, glucose and lactate can be converted to pyruvate and mixed together. In the source tissue,
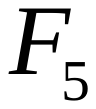
 and
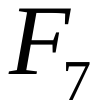
 describes the fluxes that convert glucose and lactate to pyruvate, respectively. Therefore, the predicted MID of pyruvate in the source tissue that comes from lactate and glucose can be formulated (Schematic 4).

**Schematic** **4**. Diagram of MID prediction. MIDs of substrate metabolites
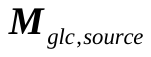
 and
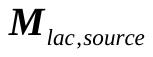
 are converted to MID of pyruvate by
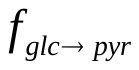
 and
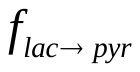
 respectively. The final MID of pyruvate is predicted based on mixture of two sources, proportional to fluxes
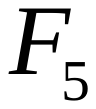
 and
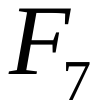
.

**Lactate in source tissue**

**Glucose in source tissue**

**Pyruvate from lactate in source tissue**

**Pyruvate from glucose in source tissue**

**Pyruvate mixing in source tissue**

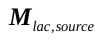

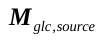

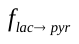

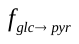

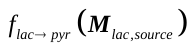

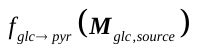

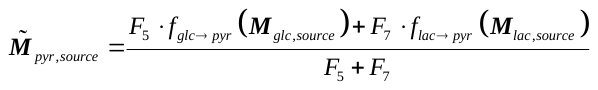

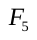

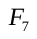

For the MID conversion function
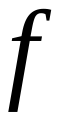
, there are three types of conversions:

(1) Transport of metabolites between plasma and tissue, such as metabolite
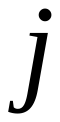
 being glucose in the plasma and metabolite
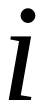
 being glucose in the source tissue. This conversion does not change the MID. Therefore,
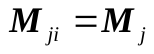
.

(2) Conversion between lactate and pyruvate, such as metabolite
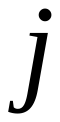
 being lactate in the source tissue and metabolite
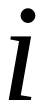
 being pyruvate in the source tissue. Because they have similar structure, conversion between lactate and pyruvate does not change MID. Therefore,
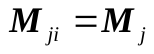
 is also valid in this category.

(3) Conversion between glucose and pyruvate, such as metabolite
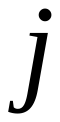
 being glucose in the source tissue and metabolite
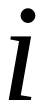
 being pyruvate in the source tissue. Because they have a different carbon number, this kind of conversion is complicated. Two special functions are designed to calculate the corresponding MIDs:

a. To calculate the MID of glucose produced by pyruvate through gluconeogenesis, a convolution is used. Suppose that the MID of pyruvate is
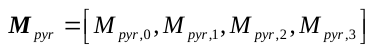
, the MID vector of glucose synthetized from pyruvate could be expressed as a convolution function:

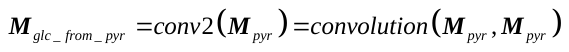
(S3)

where the discrete convolution function is defined as:

If two vectors
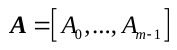
 and
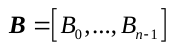
,

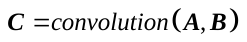
, in which:

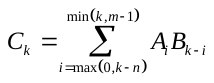
,
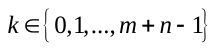
(S4)

b. To calculate the MID of pyruvate produced by glucose through glycolysis, an approximation method is used here. Suppose that the MID of glucose is
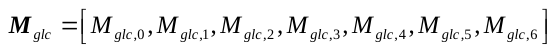
, because for glucose, carbon atoms are either all ^13^C or all ^12^C,
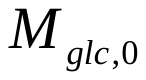
 and
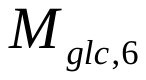
 will be dominant in MID vectors. Therefore, the MID of pyruvate from glucose could be expressed as a split function:

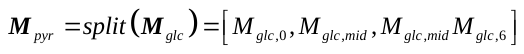
(S5)

in which
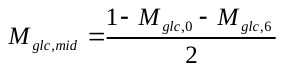

The MID of unlabeled glucose is set to
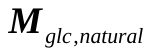
, which is a binomial distribution based on the natural abundance of ^13^C in glucose, that is:

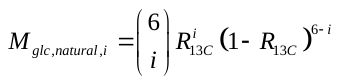
,
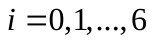
(S6)

in which
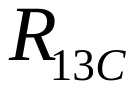
 is natural abundance of ^13^C. The MID of infused labeled glucose (substrate of
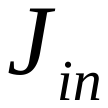
 flux in some models) is set to
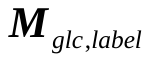
, in which all carbon atoms are ^13^C.

#### MID fitting and flux solutions

The flux solution to a MID data is obtained by minimizing the difference between predicted and experimental MID data. The difference between the predicted MID
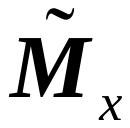
 and the experimental MID
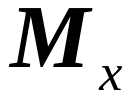
 for a metabolite
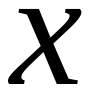
 can be defined by the Kullback–Leibler divergence
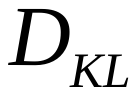
 (Kullback and Leibler, 1951), which is referred as cost function
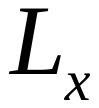
:

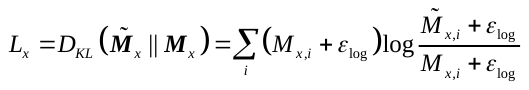
(S7)

in which
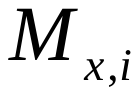
 and
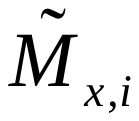
 are element
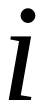
 in vector
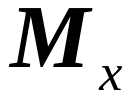
 and
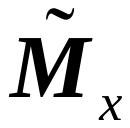
, respectively.
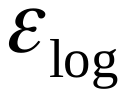
 is a small number added to maintain numerical stability.

The total cost function of a model is the sum of cost function values for selected metabolites (referred as target metabolites), which is defined as:

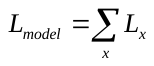
(S8)

Target metabolites in most models consisting of glucose, pyruvate and lactate in source and sink tissues. Some models may include target metabolites in plasma for better fitting.

Because each
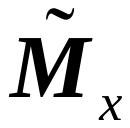
 is a function of the flux vector
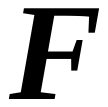
, the cost function of the model
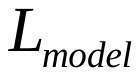
 is also a function of
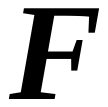
. Therefore, the flux solution can be written as:

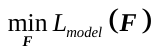
, s.t.
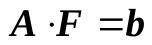
 and
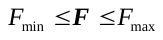
(S9)

in which
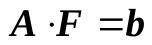
 represents the flux balance requirement and other constraints. An additional constraint of a flux range is also incorporated.

Eq. S9 represents an optimization problem with a nonlinear objective function, linear equality and inequality constraints. Therefore, it is a constrained nonlinear optimization problem. In this study, we solve this problem by sequential quadratic programming (SQP) implemented in the SciPy package (Kraft, 1988).

Similar with other iterative optimization algorithms, this algorithm starts with an initial solution and iterates to find the locally optimal point. The initial solution is generated by a linear programming (LP) problem:

, s.t. and	(S10)

in which is a uniformly distributed random vector in range [-0.4, 0.6] with the same size as . This linear programming problem is solved by a simplex algorithm implemented in SciPy package (Dantzig, 2016; Winston et al., 2003).

Because SQP can only calculate a local optimum, the LP step is repeated times to generate multiple different initial values. These initial values are fed into the SQP step to fit the flux vector respectively, and the flux vector with lowest objective value is then chosen as the final result.

Random solutions are generated by Linear Programming (eq. S10) as previously described. The objective function value is computed for each random solution. To evaluate the difference in objective function values between the computed solutions and the random solutions, a p-value is calculated from a nonparametric Wilcoxon rank-sum test implemented in the SciPy package.

#### Glucose contribution calculation

After fitting a set of fluxes from the MID data, we can use the results to calculate the relative contribution of different nutrients to the TCA cycle. For all models in this study, the contribution from each nutrient in each tissue can be regarded as one non-negative contribution flux, referred as , , etc. From those contribution fluxes, the total contribution from one metabolite is defined as:

, may not exist in some models.	(S11)

To calculate the non-negative contribution fluxes from raw flux result, we calculate all net fluxes that are connected to TCA cycle. Suppose those net fluxes are , and , they could be positive or negative. We calculate the contribution fluxes from the following formulae:

(S12)

(S13)

For each metabolite :	(S14)

For example, suppose in a tissue, glucose, lactate and pyruvate can contribute to the TCA cycle. The raw fluxes are first converted to net fluxes , and . Then, the non-negative absolute contribution fluxes , and are calculated based on eq. S14. Finally, the normalized contribution ratio can be calculated from S11 (Schematic 5).

In the simple situation with only two metabolites (lactate and glucose), eq. S14 simplifies:

If : and

If : and

If : and	(S15)

**TCA**

GLC

PYR

LAC

PYR

$$F6$$

$$F5$$

$$F7$$

$$F8$$

$$F_{11}$$

$$F9$$

$$F10$$

Raw fluxes in a single tissue model

$$F_{net,glc}=F_{5}-F_{6}>0$$

$$F_{net,lac}=F_{7}-F_{8}>0$$

$$F_{net,pyr}=F_{9}-F_{10}<0$$

**TCA**

GLC

PYR

LAC

PYR

$$F_{net,lac}$$

$$F_{11}$$

$$F_{net,glc}$$

$$F_{net,pyr}$$

**TCA**

GLC

LAC

$$F_{lac}$$

$$F_{glc}$$

**TCA**

GLC

LAC

$$R_{lac}$$

$$R_{glc}$$

$$F_{total,in}=F_{net,glc}+F_{net,lac}$$

$$F_{total,out}=-F_{net,pyr}$$

$$F_{glc}=F_{net,glc}-\frac{F_{total,out}}{F_{total,in}}F_{net,glc}$$

$$F_{lac}=F_{net,lac}-\frac{F_{total,out}}{F_{total,in}}F_{net,lac}$$

$$F_{pyr}=0$$

**(a)**

**(b)**

**(c)**

**(d)**

$$R_{glc}=\frac{F_{glc}}{F_{glc}+F_{lac}}$$

$$R_{lac}=\frac{F_{lac}}{F_{glc}+F_{lac}}$$

**Schematic 5**. Calculation of the contribution nutrients to the TCA cycle. (a) Suppose there is a one-tissue model, glucose, lactate and pyruvate can all contribute to TCA cycle. (b) Raw fluxes are first converted to net fluxes, which could be positive or negative. (c) Net fluxes are converted to contribution fluxes, which are non-negative. (d) Contribution fluxes are normalized to the calculate contribution ratio for each metabolite.

#### Solution space sampling

The dimension of the solution space in a model is calculated based on:

(S16)

in which is the number of flux variable, is the number of flux balance equations (number of eq. S1), is the number of flux constraint equations, and is the number of MID equations used to fit the model (number of eq. S7 and also elements in eq. S8). equals to 2 in basic models (model A and model B) and is larger in the complicated model.

determines the degree of freedom in the solution space. To sample the solution space uniformly, values of some fluxes are fixed during fitting (or be constant in random unfitted solutions), and the number of fixed fluxes equals to . Those fluxes with fixed values are called “free fluxes”. Those free fluxes make the dimension of the solution space of the optimization problem the same as the number of target metabolites, which prevents the problem from being overdetermined or underdetermined. The values of free fluxes are added to flux constraints equation during the optimization process.

Free fluxes are chosen based on model structure. In most models with equal to 2, fluxes and are chosen as free fluxes. The values of free fluxes are sampled uniformly in their defined ranges. When there are only two free fluxes, the whole solution space is scanned based on a lattice with discrete values on each edge (totally points) (Schematic 6(a)). As its dimension of free fluxes increases, computational cost for thorough scanning grows exponentially. Therefore, in some models with higher , we choose points with equal intervals on the diagonal of the solution space and shuffle their coordinates to cover the whole space (Schematic 6(b)).

Free flux 1

Free flux 2

Free flux 1

Free flux 2

Free flux 1

Free flux 2

**(a)**

**(b)**

**Shuffle**

**Schematic 6**. Two sampling strategies on a two dimensional flux space. (a) Uniformly scan the whole space when the dimension of free fluxes is small. (b) In high-dimension space, points on diagonal are selected uniformly, then their coordinates are shuffled to generate sample of free fluxes.

For each point of free fluxes, the LP problem is solved to obtain the initial solution. If no solution exists under the current free flux combination, this point will be discarded in the following calculation. After generating an initial solution, for the random unfitted solutions, it is directly returned for following analysis. For fitted solutions, the SQP algorithm is executed to obtain the final flux vector that minimizes the objective function (eq. S8) and the corresponding objective value . To be regarded as a feasible solution, must satisfy a series of requirements: First, should meet minimal requirement for the value of a TCA flux, which means that one or multiple TCA fluxes must be larger than a threshold . Secondly, should be small enough, which means the predicted MID data based on is close enough to experimental data. Therefore, we require the objective value must be smaller than a threshold of objective function . Only feasible solutions will be used to calculate the final distribution of glucose contribution. Therefore, the general procedure of glucose contribution analysis is shown as follows:

1. Choose the free fluxes based on the model.

2. Generate the sample of free fluxes in the solution space.

3. Optimize the objective function and solve for the corresponding flux values based on each free flux sample. Select those solutions with large enough TCA fluxes and small enough objective value.

5. Calculate glucose contribution for all feasible solutions.

5. Plot distribution of glucose contribution.

#### Parameter sensitivity

For a parameter sensitivity analysis, experimental MID data or those flux constraints are varied based on a Gaussian distribution to generate different sets. For each perturbed parameter set, a solution space sampling is executed, similar with other model, to calculate the distribution of glucose contribution. For each MID data perturbation, each experimental MID vector are multiplied with a random vector, which consists of variables with identical independent distributions, to generate raw new vector , that is:

, in which is i.i.d. and is -th element in	(S17)

follows the truncated Gaussian distribution in the range of and . Then the new raw MID vector is normalized to generate the final perturbed MID vector :

, in which is -th element in .	(S18)

The MID vector of different metabolites is multiplied by different random vector in one data perturbation. The perturbed data are used for the following glucose contribution analysis. Perturbation of other constraints is similar with MID data perturbation. In each perturbation, the target constraint is multiplied by a random variable, that is:

(S19)

(S20)

(S21)

Similar with the MID data, follows the truncated Gaussian distribution in the range of or . The perturbed parameters are used for the following glucose contribution analysis.

#### Hypoxia correction

Tissue extraction introduces issues due to hypoxia. To estimate these effects to the final glucose contribution, a correction to the MID data is introduced to simulate this process. The hypoxic state includes two major events: glycogen breakdown in source tissue and elevated lactate generation in sink tissue (figure S3A). To correct for the effect of hypoxia, the current data is assumed to be measured under hypoxia, which means the current MID of metabolites is a mixture of that metabolite in the original tissue and product of activated reaction under the hypoxia state. Specifically, the MID of glucose in the source tissue is mixture of (80%) original glucose MID and (20%) hydrolyzed glucose from glycogen (unlabeled MID), and the MID of lactate in the sink tissue is mixture of (80%) original one and (20%) reductive product from pyruvate (same MID as pyruvate in sink tissue) (figure S3B). From this assumption we calculate putative original MIDs of these two metabolites. If there is any negative item in MID, assign all of them to and re-nomalized each MID to ensure sum of them equals to 1. Use those processed MID to do the same fitting and calculation of glucose contribution as Model A.

#### Ternary graph plotting

In those models with three circulating metabolites, glucose, pyruvate and lactate can all contribute to the contribution to the TCA cycle. Therefore, the ternary graph is plotted to display the distribution of their relative contribution ratio in one figure. The ternary graphs are plotted using a python package, python-ternary (<https://github.com/marcharper/python-ternary>).

For each free flux sample, the contribution from glucose , from pyruvate and from lactate are calculated based on eq. S14. Each triple set in ternary space corresponds to the contribution of one sample of a free flux set with objective value lower than threshold. To better display the distribution of contribution the set of three fluxes, those points are binned and used to make the density heatmap. as figure 5D and E. Because of the limitation of ternary plot package, a complicated protocol is designed to reflect the point density (Schematic 8).

First, those sets in ternary space are transformed to the Cartesian coordinate system in the space by the following equations:

(S22)

The contribution of triplet set of all solution points are mapped onto the Cartesian system and binned in a two dimensional (2D) grid with bins on each edge. The output matrix is a square matrix with items. A Gaussian kernel matrix with the same size as from a two dimensional Gaussian distribution with the center at origin and covariance matrix as is constructed. Then, the binned contribution matrix and kernel matrix are convoluted to obtain the final density matrix based on a 2D discrete convolution rule:

If and are square matrices with m- and n-dimension respectively,

in which . (Fill with 0 if out of scope)	(S23)

In the final ternary graph, the triangle is divided into smaller hexagons. For each hexagon, its center coordinates in three-dimensional space are mapped to 2D Cartesian space to get based on eq. S22. This hexagon is colored based on the interpolated onto the density matrix .

#### Software implementation

Scripts in this study are implemented by Python 3.6. Results are running on a desktop PC with an i7-8700 CPU. To reduce the running time, some strategies such as parallel based processing are utilized. Each model requires around 10 ~ 50 hours of CPU running time.

#### Common parameters:

| Parameter | Comment | Value |
| --- | --- | --- |
|  | Natural abundance of ^13^C | 0.01109 |
|  | MID of labeled infusion glucose |  |
|  | Small number to increase numeric stability in log function | 1e-10 |
|  | Small number to increase numeric stability in MID normalization | 1e-5 |

$$\boldsymbol{R}_{\boldsymbol{glu}}$$

$$\boldsymbol{R}_{\boldsymbol{lac}}$$

$$\boldsymbol{R}_{\boldsymbol{pyr}}$$

0.0

1.0

0.2

0.4

1.0

0.6

0.8

1.0

0.0

0.2

0.4

0.6

0.8

0.0

0.2

0.4

0.6

0.8

**Coordinate transformation**

0.0

0.2

0.4

0.6

0.8

1.0

0.0

0.2

0.4

0.6

0.8

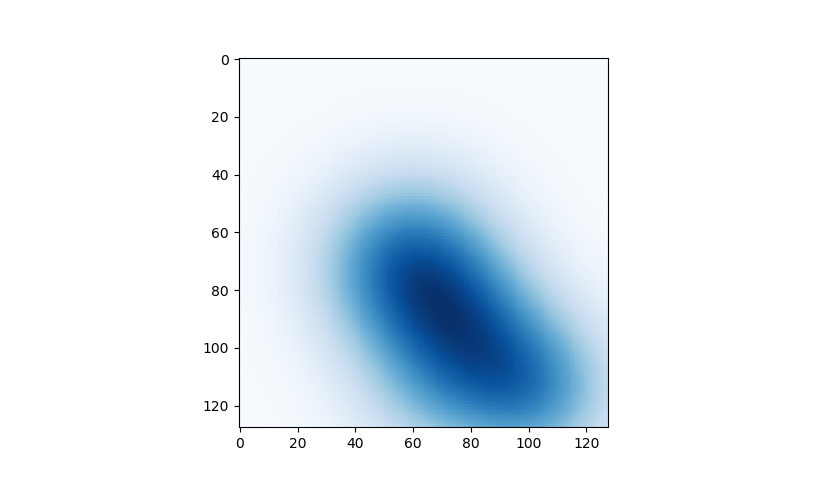

**Convolution with Gaussian distribution**

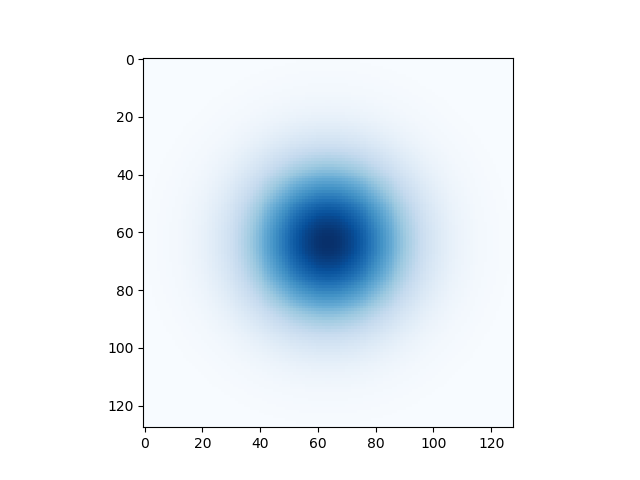

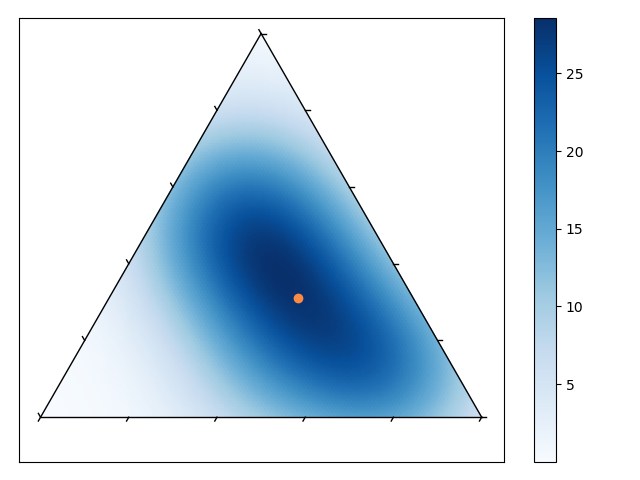

$$\boldsymbol{R}_{\boldsymbol{glu}}$$

$$\boldsymbol{R}_{\boldsymbol{lac}}$$

$$\boldsymbol{R}_{\boldsymbol{pyr}}$$

0.0

1.0

0.2

0.4

1.0

0.6

0.8

1.0

0.0

0.2

0.4

0.6

0.8

0.0

0.2

0.4

0.6

0.8

**Mean point**

**Coordinate transformation**

**Schematic 7**. The density distribution of contribution from glucose, lactate and pyruvate. The contribution triple set is first transformed to Cartesian system , and convolved with a Gaussian kernel. Then, the resulting distribution is mapped back to ternary system with mean value calculated.

### Specific models

Based on general protocols described above, many models are implemented in this study. They are different in data source and metabolites, tissues and parameters included in those models. Relationships between these models are shown in Schematic 8.

**Schematic 8**. Relationships between different models. Model A is the basic model. Model C is the model A with one more sink tissue. Model B is the model A fitted with high infusion data. Model D is the model A with one more circulating metabolite (pyruvate), while model E is the model D fitted with high infusion data.

Model A

Model B

Model C

Model D

Model E

High infusion data

High infusion data

Add one more sink tissue

Add one more circulating metabolite

Add one more circulating metabolite

**Basic model**

#### Model A: basic model for two tissues (figure 1, S1, 2, S2)

**Flux balance equations:**

Glucose in source tissue:

Pyruvate in source tissue:

Lactate in source tissue:

Glucose in plasma:

Lactate in plasma:

Glucose in sink tissue:

Pyruvate in sink tissue:

Lactate in sink tissue:

**Flux constraints:**

Supplement glucose flux:

Glucose turnover flux:

Lactate turnover flux:

**MID data:**

Glucose in source tissue:

Pyruvate in source tissue:

Lactate in source tissue:

Glucose in plasma:

Lactate in plasma:

Glucose in sink tissue:

Pyruvate in sink tissue:

Lactate in sink tissue:

**MID predictions:**

Glucose in source tissue:

Pyruvate in source tissue:

Lactate in source tissue:

Glucose in sink tissue:

Pyruvate in sink tissue:

Lactate in sink tissue:

**Cost function:**

(S24)

**Glucose contribution calculation:**

After fitting a result , glucose contribution is calculated based on eq. S11 and S15. We first calculate , , and :

, , ,	(S25)

Therefore, and can be calculated by:

If and , ,

If and , ,

If and , ,	(S26)

Because and it must be non-negative, it is impossible that and are both negative.

The and in the sink tissue have a similar form by replacing to in eq. S26.

Therefore, the glucose contribution of sink tissue and in complete model can be calculated as:

(S27)

(S28)

Similarly, the lactate contribution can also be calculated as:

(S29)

(S30)

**Free fluxes and sampling:**

and are chosen as free fluxes. Because of limitation of circulatory flux of glucose, the common upper bound for them is . Each of them is uniformly sampled from for different values. Therefore, the total sample number is . For each sampled point, if or after optimization, this sample is filtered.

**Data source:**

The data to fit this model is the low-infusion data set. The source tissue is liver, while the sink tissue is one from heart, brain, skeletal muscle, kidney, lung, pancreas, small intestine and spleen, respectively. If not mentioned, MID data from glucose-infused M1 is used by default. Glucose-infused M5 and M9, and lactate-infused M3, M4, M10 and M11 are also analyzed to prove the data robustness.

**Parameter table:**

| Category | Parameter | Comment | Value |
| --- | --- | --- | --- |
| Model |  | Total flux number | 19 |
|  |  | Number of flux balance equations | 8 |
|  |  | Number of flux constraints (not including free fluxes) | 3 |
|  |  | Number of MID predictions | 6 |
|  |  | Number of free fluxes | 2 |
|  |  | Minimal flux value | 1 |
|  |  | Maximal flux value | 500 |
|  |  | Value of supplement glucose flux in source tissue | 35 |
|  |  | Value of glucose turnover flux | 150.9 |
|  |  | Value of lactate turnover flux | 374.4 |
| Optimization |  | Repeat number to optimize the cost function | 10 |
|  |  | Objective value threshold to accept the fitting result | 0.1 |
| Sample |  | Sample number for each free flux | 1000 |
|  |  | Minimal TCA flux value | 2 |

#### Distribution of glucose contribution

The contribution ratio $R_{glc}$ only reflects contribution ratios of circulating metabolites in sink tissue, and consequently their distributions are significantly different from $R_{glc}^{'}$. $R_{glc}$ usually shows a bimodal distribution, and most of them concentrate around 1. This kind of special distribution can be explained by model structure:

In the two-tissue model, flux balance requirement only allow three patterns for net fluxes (Schematic 7). Among this three patterns, the pattern with contribution ratio $0<R_{glc}<1$ has smaller solution space than other two patterns, and solutions with this pattern are evenly distributed between the range 0 to 1. Therefore, in violin plots, $R_{glc}$ of most feasible solutions have bimodal distributions with $R_{glc}=0$ or $1$. The distribution that in most cases $R_{glc}$ concentrates on 1 shows net fluxes follow the pattern of Cori cycle, in which in sink tissue glucose is transformed to lactate and in source tissue lactate is transformed to pyruvate or glucose (Schematic 7). Therefore, these results are consistent with the conclusion that circulating glucose is the major contribution to TCA cycle in sink tissue. Situations are also similar for other complicated models.

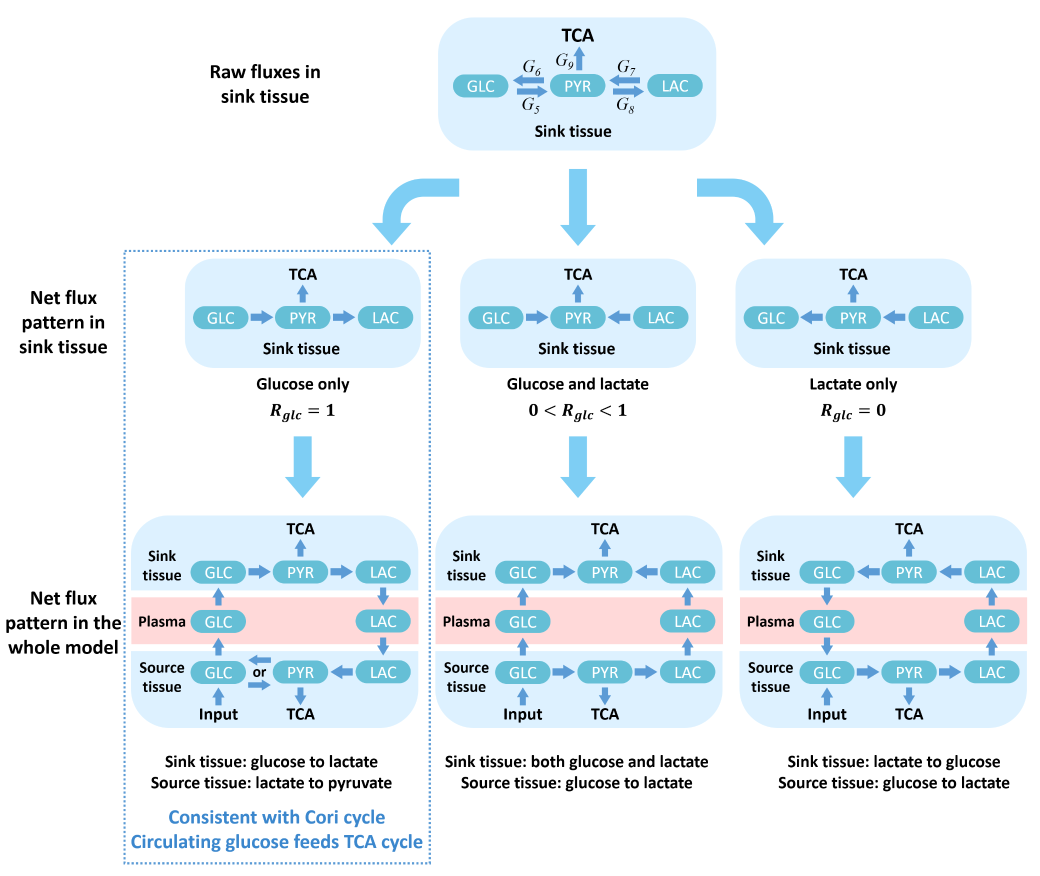

**Schematic 7**. Three possible patterns of feasible solutions in our model, which correspond to three glucose contribution ratios: $R_{glc}=1$, $0<R_{glc}<1$ or $R_{glc}=0$. In the first pattern, glucose is transformed to lactate in sink tissue and lactate is transformed to pyruvate or glucose, which is consistent with Cori cycle. In our results $R_{glc}$ concentrates on 1 supports our conclusion that circulating glucose feeds TCA cycle in most kinds of sink tissue.

#### Parameter sensitivity for model A (figure 3)

MID data and three parameters are perturbed individually and used for analysis based on model A. All model constructions and unperturbed parameters are also same as model A. Only the resolution to sample the solution space is reduced to increase efficiency.

**Data source:**

Similar with model A, this part uses the low-infusion data set. In all perturbations, the source tissue is liver and the sink tissue is heart. Only the MID data from M1 is used.

**Parameter table:**

(Underlined items indicate differences from those in model A)

| Category | Parameter | Comment | Value |
| --- | --- | --- | --- |
| Model |  | Total flux number | 19 |
|  |  | Number of flux balance equations | 8 |
|  |  | Number of flux constraints (not including free fluxes) | 3 |
|  |  | Number of MID predictions | 6 |
|  |  | Number of free fluxes | 2 |
|  |  | Minimal flux value | 1 |
|  |  | Maximal flux value | 1000 |
|  |  | Initial value of supplement glucose flux in source tissue | 100 |
|  |  | Initial value of glucose turnover flux | 150.9 |
|  |  | Initial value of lactate turnover flux | 374.4 |
| Optimization |  | Repeat number to optimize the cost function | 10 |
|  |  | Objective value threshold to accept the fitting result | 0.2 |
| Sample |  | Sample number for each free flux | 100 |
|  |  | Minimal TCA flux value | 2 |
| Parameter sensitivity |  | Variance of perturbation random variable for MID data | 0.5 |
|  |  | Variance of perturbation random variable for constant fluxes | 0.2 |
|  |  | Variance range of MID data | $\pm\left[ 0.1, 0.9 \right]$ |
|  |  | Variance range of constant fluxes | $\pm\left[ 0.05, 0.6 \right]$ |
|  |  | Number of different perturbations generated for sensitivity analysis | 100 |

#### Hypoxia correction for model A (figure S3)

MID data are corrected and used for analysis based on model A. All model constructions and unperturbed parameters are also same as model A.

**Data source:**

Similar with model A, this part uses the low-infusion data set. The source tissue is liver and the sink tissue is heart. Only the MID data from M1 is used.

**Parameter table:**

(Underlined items indicate differences from those in model A)

| Category | Parameter | Comment | Value |
| --- | --- | --- | --- |
| Model |  | Total flux number | 19 |
|  |  | Number of flux balance equations | 8 |
|  |  | Number of flux constraints (not including free fluxes) | 3 |
|  |  | Number of MID predictions | 6 |
|  |  | Number of free fluxes | 2 |
|  |  | Minimal flux value | 1 |
|  |  | Maximal flux value | 500 |
|  |  | Value of supplement glucose flux in source tissue | 35 |
|  |  | Value of glucose turnover flux | 150.9 |
|  |  | Value of lactate turnover flux | 374.4 |
| Optimization |  | Repeat number to optimize the cost function | 10 |
|  |  | Objective value threshold to accept the fitting result | 0.1 |
| Sample |  | Sample number for each free flux | 1000 |
|  |  | Minimal TCA flux value | 2 |
| Hypoxia correction |  | Assumed mixture ratio for hypoxia correction | 20% |

#### Model B: model for high-infusion data (figure 4, S4)

(Underlined items indicate differences from those in model A)

**Flux balance equations:**

Glucose in source tissue:

Pyruvate in source tissue:

Lactate in source tissue:

Glucose in plasma:

Lactate in plasma:

Glucose in sink tissue:

Pyruvate in sink tissue:

Lactate in sink tissue:

**Flux constraints:**

This model removes the glucose turnover flux constraint. Alternatively, it adds glucose in plasma to target metabolites.

Supplement glucose flux:

Infusion glucose flux:

Lactate turnover flux:

**MID data:**

Glucose in source tissue:

Pyruvate in source tissue:

Lactate in source tissue:

Glucose in plasma:

Lactate in plasma:

Glucose in sink tissue:

Pyruvate in sink tissue:

Lactate in sink tissue:

**MID predictions:**

Glucose in source tissue:

Pyruvate in source tissue:

Lactate in source tissue:

Glucose in plasma:

Glucose in sink tissue:

Pyruvate in sink tissue:

Lactate in sink tissue:

**Cost function:**

(S31)

**Glucose contribution calculation:**

Glucose contribution calculation in this model is same as model A. The raw flux result is processed by eq. S25 and S26 to calculate glucose and lactate contribution fluxes. Finally, Eqs. S27 and S29 are utilized to calculate the relative glucose and lactate contribution and , while Eqs. S28 and S30 are for and .

**Free fluxes and sampling:**

in this model is still 2. Free fluxes are also and . This model removes constraint on the glucose turnover flux, and thus the free fluxes have a wider range. Each flux is uniformly sampled from for different values. Therefore, the total sample size is still . For each sampled point, if or after optimization, this sample is filtered out.

**Data source:**

The data to fit this model is the high-infusion data set. The source tissue is liver, while the sink tissue is skeletal muscle. The MID data from mouse M1, M2, M3 and M4 are used.

**Parameter table:**

Because of the higher infusion flux, the glucose turnover flux in plasma will increase, and thus lactate turnover flux should also increase. Furthermore, the higher labeling ratio decreases fitting accuracy (fig. S4). Therefore, the threshold of objective function was also increased.

| Category | Parameter | Comment | Value |
| --- | --- | --- | --- |
| Model |  | Total flux number | 20 |
|  |  | Number of flux balance equations | 8 |
|  |  | Number of flux constraints (not including free fluxes) | 3 |
|  |  | Number of MID predictions | 7 |
|  |  | Number of free fluxes | 2 |
|  |  | Minimal flux value | 1 |
|  |  | Maximal flux value | 1000 |
|  |  | Value of supplement glucose flux in source tissue | 80 |
|  |  | Value of glucose infusion flux | 111.1 |
|  |  | Value of lactate turnover flux | 400 |
| Optimization |  | Repeat number to optimize the cost function | 10 |
|  |  | Objective value threshold to accept the fitting result | 0.25 |
| Sample |  | Sample number for each free flux | 1500 |
|  |  | Maximal flux value of two free fluxes | 300 |
|  |  | Minimal TCA flux value | 2 |

#### Model C: model for three tissues (figure 5, S5)

(Underlined items indicate differences from those in model A)

**Flux balance equations:**

Glucose in source tissue:

Pyruvate in source tissue:

Lactate in source tissue:

Glucose in sink tissue 1:

Pyruvate in sink tissue 1:

Lactate in sink tissue 1:

Glucose in sink tissue 2:

Pyruvate in sink tissue 2:

Lactate in sink tissue 2:

Glucose in plasma:

Lactate in plasma:

**Flux constraints:**

Supplement glucose flux:

Glucose turnover flux:

Lactate turnover flux:

**MID data:**

Glucose in source tissue:

Pyruvate in source tissue:

Lactate in source tissue:

Glucose in sink tissue 1:

Pyruvate in sink tissue 1:

Lactate in sink tissue 1:

Glucose in sink tissue 2:

Pyruvate in sink tissue 2:

Lactate in sink tissue 2:

Glucose in plasma:

Lactate in plasma:

**MID predictions:**

Glucose in source tissue:

Pyruvate in source tissue:

Lactate in source tissue:

Glucose in sink tissue 1:

Pyruvate in sink tissue 1:

Lactate in sink tissue 1:

Glucose in sink tissue 2:

Pyruvate in sink tissue 2:

Lactate in sink tissue 2:

**Cost function:**

(S32)

**Glucose contribution calculation:**

Slightly different from that in model A, after fitting a result , , , , , and can be calculated from raw flux values:

(S33)

Therefore, , , , , and in different tissue can be calculated based on eq. S26. Then, the glucose contribution in sink tissue and in complete model can be calculated as:

(S34)

(S35)

Similarly, the lactate contribution and can also be calculated as:

(S36)

(S37)

**Free fluxes and sampling:**

in this model is 5. Therefore, , , , and are chosen as free fluxes. These fluxes are constrained by circulatory fluxes of glucose and lactate (see glucose turnover flux and lactate turnover flux in “flux constraints” section). Therefore, their maximal value is bounded by or . Specifically, dynamic ranges of , and are , and those of and are . Those dynamic ranges constitute a 5-dimension solution space . To uniformly sample in , we pick points from its diagonal and shuffle the five coordinates of those points. Those sample points are used for following analysis. For each sampled point, if and after optimization, this sample is filtered.

**Data source:**

The data to fit this model is the low-infusion data set. The source tissue is liver, the sink tissue 1 and sink tissue 2 are combinations from heart, brain and skeletal muscle. MID data from mouse M1 is used.

**Parameter table:**

Because the cost function includes more MID data during the optimization process, the threshold of objective function also increases. The sample number is set to be close to the previous total sample number .

| Category | Parameter | Comment | Value |
| --- | --- | --- | --- |
| Model |  | Total flux number | 28 |
|  |  | Number of flux balance equations | 11 |
|  |  | Number of flux constraints (not including free fluxes) | 3 |
|  |  | Number of MID predictions | 9 |
|  |  | Number of free fluxes | 5 |
|  |  | Minimal flux value | 1 |
|  |  | Maximal flux value | 700 |
|  |  | Value of supplement glucose flux in source tissue | 40 |
|  |  | Value of glucose turnover flux | 150.9 |
|  |  | Value of lactate turnover flux | 374.4 |
| Optimization |  | Repeat number to optimize the cost function | 10 |
|  |  | Objective value threshold to accept the fitting result | 0.15 |
| Sample |  | Total sample number in solution space | 3×10^6^ |
|  |  | Minimal TCA flux value | 2 |

#### Model D: model for three circulating metabolites for low-infusion data (figure 6B, D, S6A, C, E, G, I)

(Underlined items indicate differences from those in model A)

**Flux balance equations:**

Glucose in source tissue:

Pyruvate in source tissue:

Lactate in source tissue:

Glucose in plasma:

Lactate in plasma:

Pyruvate in plasma:

Glucose in sink tissue:

Pyruvate in sink tissue:

Lactate in sink tissue:

**Flux constraints:**

Supplement glucose flux:

Glucose turnover flux:

Lactate turnover flux:

Pyruvate turnover flux:

**MID data:**

Glucose in source tissue:

Pyruvate in source tissue:

Lactate in source tissue:

Glucose in plasma:

Lactate in plasma:

Pyruvate in plasma:

Glucose in sink tissue:

Pyruvate in sink tissue:

Lactate in sink tissue:

**MID predictions:**

Glucose in source tissue:

Pyruvate in source tissue:

Lactate in source tissue:

Glucose in sink tissue:

Pyruvate in sink tissue:

Lactate in sink tissue:

Lactate in plasma:

Pyruvate in plasma:

**Cost function:**

(S38)

**Glucose contribution calculation:**

Because there are three nutrients that contribute to the TCA cycle, after fitting a result , the glucose, lactate and pyruvate contribution ratio, , and respectively, are calculated based on eq. S11.

Firstly, the net fluxes connected to the TCA cycle can be calculated:

(S39)

The total in and out fluxes for the TCA cycle in the source tissue ( and ) and in the sink tissue ( and ) can be calculated based on eq. S12, S13 and those net fluxes in eq. S39. Contribution fluxes of glucose , lactate and pyruvate in the source tissue can be calculated from eq. S14 and net fluxes in eq. S39. Similarly, , and in the sink tissue can also be calculated. Therefore, the contribution ratio from three metabolites can be calculated as:

(S40)

(S41)

(S42)

(S43)

(S44)

(S45)

**Free fluxes and sampling:**

in this model is 5. Therefore, , , , and are chosen as free fluxes. These fluxes are constrained by circulatory fluxes of glucose, lactate and pyruvate (see glucose turnover flux, lactate turnover flux and pyruvate turnover flux in “flux constraints” section). Therefore, their upper bounds are set by , or . Specifically, the dynamic ranges of and are , those of and are , and that of is . Those dynamic ranges constitute a 5-dimensional solution space . Similar with what was computed in model C, we pick points uniformly from the diagonal of and shuffle the five coordinates of those points. Those sampled points are generated. For each sampled point, if or after optimization, this sample is filtered.

**Data source:**

The data to fit this model is the low-infusion data set. Similar with model A, the source tissue is liver and the sink tissue is heart. The MID data from mouse M1 are used.

**Parameter table:**

Because this model includes more circulating metabolites, the value of input glucose flux is set slightly higher than that in model A. Similar with model C, the threshold of the objective function also increases, and the sample number is set to the same value. Parameters for the ternary graph are set to provide high resolution.

| Category | Parameter | Comment | Value |
| --- | --- | --- | --- |
| Model |  | Total flux number | 26 |
|  |  | Number of flux balance equation | 9 |
|  |  | Number of flux constraints (not including free fluxes) | 4 |
|  |  | Number of MID predictions | 8 |
|  |  | Number of free fluxes | 5 |
|  |  | Minimal flux value | 1 |
|  |  | Maximal flux value | 800 |
|  |  | Value of supplement glucose flux in source tissue | 60 |
|  |  | Value of glucose turnover flux | 150.9 |
|  |  | Value of lactate turnover flux | 374.4 |
|  |  | Value of pyruvate turnover flux | 57.3 |
| Optimization |  | Repeat number to optimize the cost function | 10 |
|  |  | Objective value threshold to accept the fitting result | 0.15 |
| Sample |  | Total sample number in solution space | 1×10^6^ |
|  |  | Minimal TCA flux value | 2 |
| Ternary graph |  | Variance of gaussian kernel in ternary graph | 0.15 |
|  |  | Resolution of ternary graph | 256 |

#### Model E: model for three circulating metabolites for high-infusion data (figure 6E, S6B, D, F, J)

(Underlined items indicate differences from those in model A)

The only difference between this model and model D is the infusion flux.

**Flux balance equations:**

Glucose in source tissue:

Pyruvate in source tissue:

Lactate in source tissue:

Glucose in plasma:

Lactate in plasma:

Pyruvate in plasma:

Glucose in sink tissue:

Pyruvate in sink tissue:

Lactate in sink tissue:

**Flux constraints:**

Supplement glucose flux:

Infusion glucose flux:

Lactate turnover flux:

Pyruvate turnover flux:

**MID data:**

Glucose in source tissue:

Pyruvate in source tissue:

Lactate in source tissue:

Glucose in plasma:

Lactate in plasma:

Pyruvate in plasma:

Glucose in sink tissue:

Pyruvate in sink tissue:

Lactate in sink tissue:

**MID predictions:**

Glucose in source tissue:

Pyruvate in source tissue:

Lactate in source tissue:

Glucose in sink tissue:

Pyruvate in sink tissue:

Lactate in sink tissue:

Glucose in plasma:

Lactate in plasma:

Pyruvate in plasma:

**Cost function:**

(S46)

**Glucose contribution calculation:**

The glucose contribution calculation in this model is same as in model D. After fitting a flux vector , the contribution to the TCA cycle from glucose , from lactate and from pyruvate in sink tissue can be calculated by eq. S40, S42 and S44, respectively. Similarly, those contribution in complete model , and can also be calculated by eq. S41, S43 and S45.

**Free fluxes and sampling:**

It is the same as in model D. , , , and are chosen as free fluxes. Dynamic ranges of and are , those of and are , and that of is . points are picked uniformly from its diagonal and the five coordinates of those points are then randomly shuffled. Those sample points are generated. For each sampled point, if or after optimization, this sample is filtered.

**Data source:**

The data to fit this model is the high-infusion data set. Similar as in model C, the source tissue is liver and the sink tissue is skeletal muscle. MID data from mouse M1 are used.

**Parameter table:**

Most parameters are same with model D. Circulatory fluxes of lactate and pyruvate are increased to adapt to the higher glucose infusion flux. Similar with model C, the higher labeling ratio decreases the fitting accuracy, hence requiring a higher tolerance threshold of objective function.

| Category | Parameter | Comment | Value |
| --- | --- | --- | --- |
| Model |  | Total flux number | 26 |
|  |  | Number of flux balance equations | 9 |
|  |  | Number of flux constraints (not including free fluxes) | 4 |
|  |  | Number of MID predictions | 8 |
|  |  | Number of free fluxes | 5 |
|  |  | Minimal flux value | 1 |
|  |  | Maximal flux value | 1000 |
|  |  | Value of supplement glucose flux in source tissue | 150 |
|  |  | Value of glucose infusion flux | 111.1 |
|  |  | Value of lactate turnover flux | 400 |
|  |  | Value of pyruvate turnover flux | 70 |
| Optimization |  | Repeat number to optimize the cost function | 10 |
|  |  | Objective value threshold to accept the fitting result | 0.4 |
| Sample |  | Total sample number in solution space | 3×10^6^ |
|  |  | Minimal TCA flux value | 2 |
| Ternary graph |  | Variance of gaussian kernel in ternary graph | 0.15 |
|  |  | Resolution of ternary graph | 256 |

### Physiological feasibility of solutions

Physiological feasibility is a key feature for biomedical models. In the results shown in main figures, solutions that have low TCA flux are considered as infeasible in physiological conditions, and therefore filtered out. Here we provide an example of a solution without this filter for comparison. For simplicity, solutions with filter (in main figures) are referred as filtered results, while the results without filter (in this section) are referred as unfiltered results.

Solving process is identical with that in main text. If not specified, all other parameters are same.

#### Model A: basic model for two tissues

| Category | Parameter | Comment | Value |
| --- | --- | --- | --- |
| Model |  | Maximal flux value | 1000 |
|  |  | Value of supplement glucose flux in source tissue | 100 |

In this model, the fitting process is identical to filtered results of Model A. Compared with filtered results in the main text, fitting precision has not been changed: MID prediction and distribution of objective function without filter are almost same as results with filter (Schematic 9a-h, j-p, compared with figure S1). Distributions of most fluxes are also similar, but F9 and G9, which are two TCA fluxes to source and sink tissue respectively, are significantly different: Before filtering, boxplot shows quantiles of F9 and G9 are closed to extreme value (Schematic 9i). This kind of distribution means one of TCA fluxes is optimized to near zero in many results, which should not occur in physiological condition. After filtering, F9 and G9 are more concentrated on half of , and two TCA fluxes are well balanced, which should be more physiologically feasible (figure S1I).

The definition of glucose contribution in the complete model $R_{glc}^{'}$ is also the same (figure S2A, S2B). Distribution of $R_{glc}^{'}$ in unfiltered results is generally similar with that in filtered ones, but the distribution is more concentrated (Schematic 10, compared with figure S2B-I. Notice that the arrangement of sink tissue and mice is different in two results). This may due to high enrichment of extreme value of TCA fluxes F9 and G9.

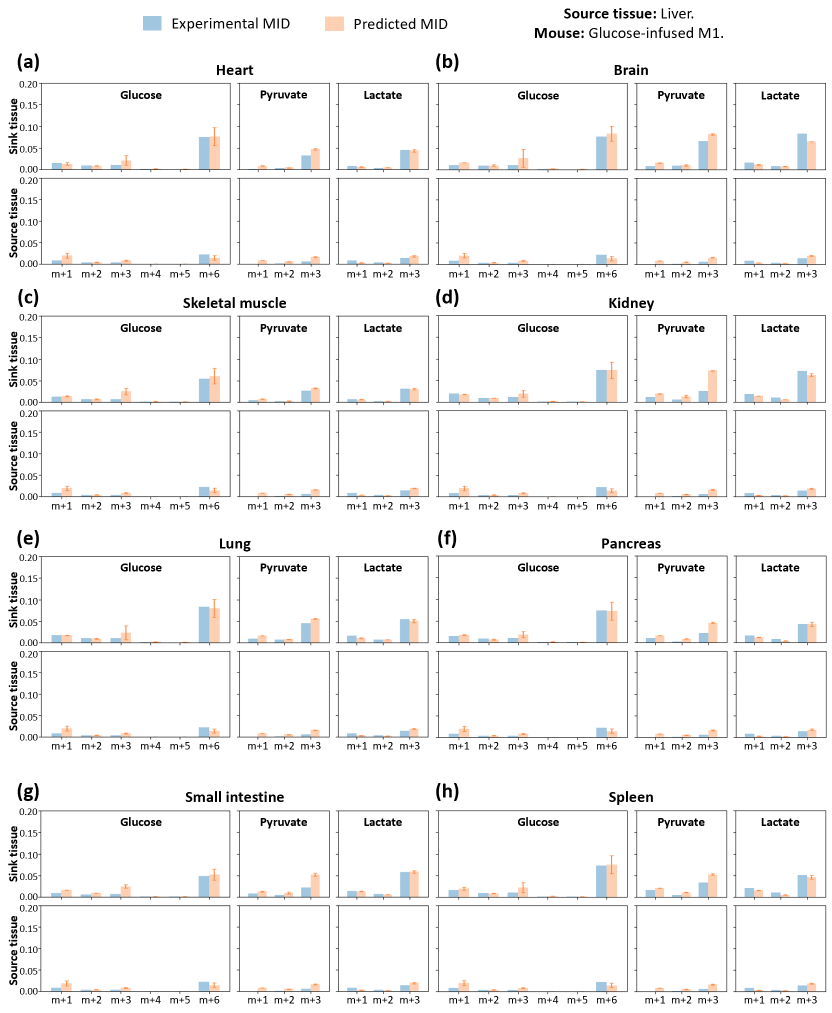

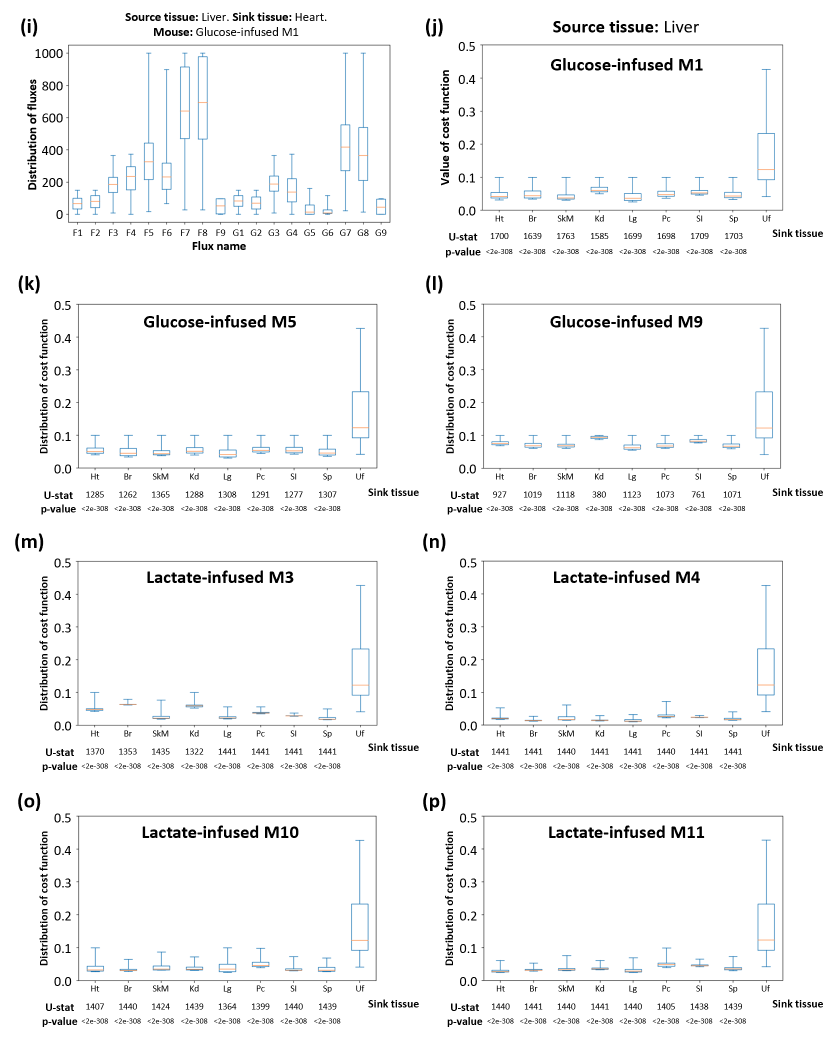

**Schematic 9**. Detailed information of model fitting. (a-h) Comparison of experimental and predicted MID in the model. Average of predicted MID in all feasible solutions are displayed. Standard deviation is also displayed as error bar. In most cases experimental MID could be well predicted by this model. Because of the low abundance, only isotopomers with more than one ^13^C are displayed. (i) Distribution of 18 variable fluxes in all feasible solutions. Source tissue is liver and sink tissue is heart. (j-p) Distribution of cost functions of feasible solutions fitted with different sink tissue and data from different mice, compared with unfitted control data. Source tissue in all fittings is liver. U-statistics of rank-sum test and p-values are displayed. Ht: heart, Br: brain, SkM: skeletal muscle, Kd: kidney, Lg: lung, Pc: pancreas, SI: small intestine, Sp: spleen, Uf: data from random unfitted control.

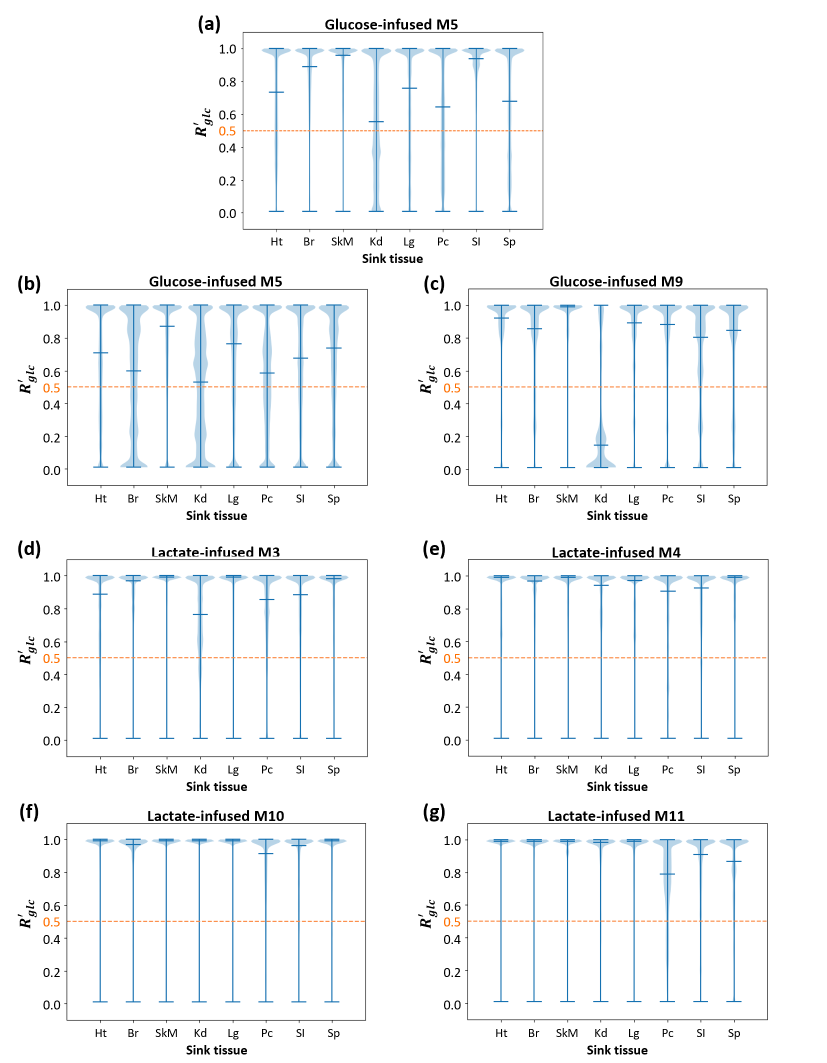

**Schematic 10**. Distribution of glucose contribution based on the model with different sink tissues fitted with different mice. For most sink tissues in all mice, the median of glucose contribution is close to or higher than 0.5, which means glucose contributes more than lactate to the TCA cycle.The orange dash line represents 0.5 threshold. Data set is from glucose-infused mice (M1, M5, M9) and lactate-infused mice (M3, M4, M10, M11) in Hui et al, 2017.

#### Parameter sensitivity for model A

| Category | Parameter | Comment | Value |
| --- | --- | --- | --- |
| Model |  | Maximal flux value | 1000 |
|  |  | Value of supplement glucose flux in source tissue | 100 |
| Parameter sensitivity |  | Variance range of constant fluxes | $\pm\left[ 0.1, 0.9 \right]$ |
|  |  | Variance of perturbation random variable for constant fluxes | 0.5 |

Glucose contribution for lactate circulatory flux in this calculation is more robust than that in main figures (Schematic 11, figure 3C). First reason is this calculation count for robustness of glucose contribution in complete model ($R_{glc}^{'}$), while that in the main figure is glucose contribution in sink tissue ($R_{glc}$). $R_{glc}$ may be more sensitive to perturbation on lactate circulatory flux. Another reason is current calculation has not been filtered, and TCA fluxes F9 and G9 concentrate on extreme value, which causes the relatively concentrated distribution of glucose contribution. This pattern may be more resistant to perturbation on parameters.

#### Model B: model for high-infusion data

| Category | Parameter | Comment | Value |
| --- | --- | --- | --- |
| Model |  | Maximal flux value | 2000 |
|  |  | Value of supplement glucose flux in source tissue | 100 |

Similarly, the fitting process except filtering is identical to filtered results of Model B. Compared with filtered results in the main text, fitting precision has not been changed: MID prediction and distribution of objective function in unfiltered results are almost same as filtered results (Schematic 12A, C, figure 4D, S4A). The distribution of fluxes for feasible solutions has significant changes: because the is higher in this calculation, G7 and G8 are significantly larger than other fluxes, which is less feasible in physiological condition (Schematic 12B, compared with figure S4B). Furthermore, similar with Model A, F9 and G9 tend to be optimized to extreme value, such as those in M1, M2 and M4 (Schematic 12B). After filtering, as espected, F9 and G9 concentrate on intermediate value (figure S4B).

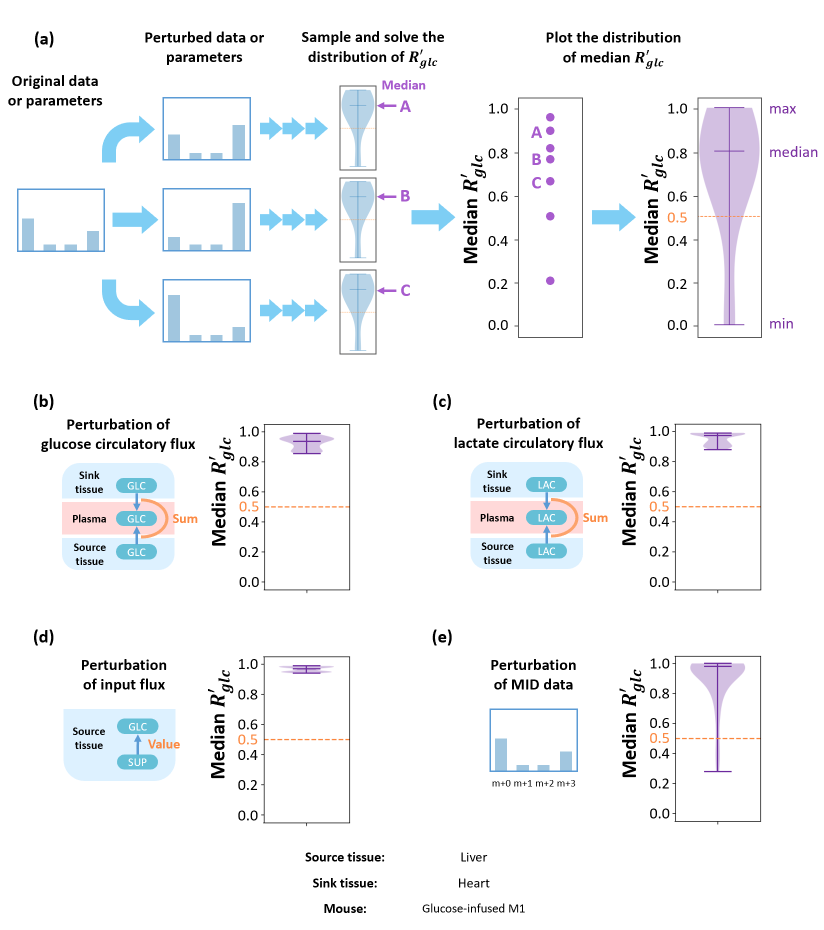

**Schematic 11**. Parameter sensitivity analysis. (a) Original MID data or constraint parameters are randomly perturbed and used in the following analysis. The resulting distribution of the glucose contribution for each perturbation is calculated, and their medians are collected. Distribution of medians reflects parameter sensitivities for this model. The distribution of medians under perturbation of glucose circulatory flux (b), lactate circulatory flux (c), input flux in source tissue (d) and MID data (e). Most of the medians are above the 0.5 threshold, which implies that under most perturbations, glucose contributes more than lactate to the TCA cycle. Data set is from glucose-infused mouse M1 in Hui et al, 2017. Source tissue is liver and sink tissue is heart.

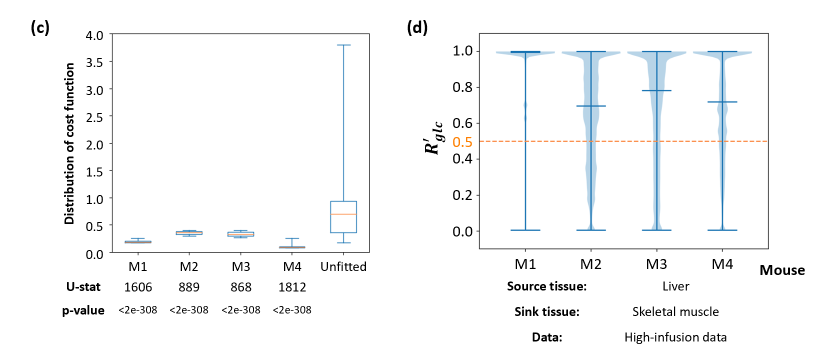

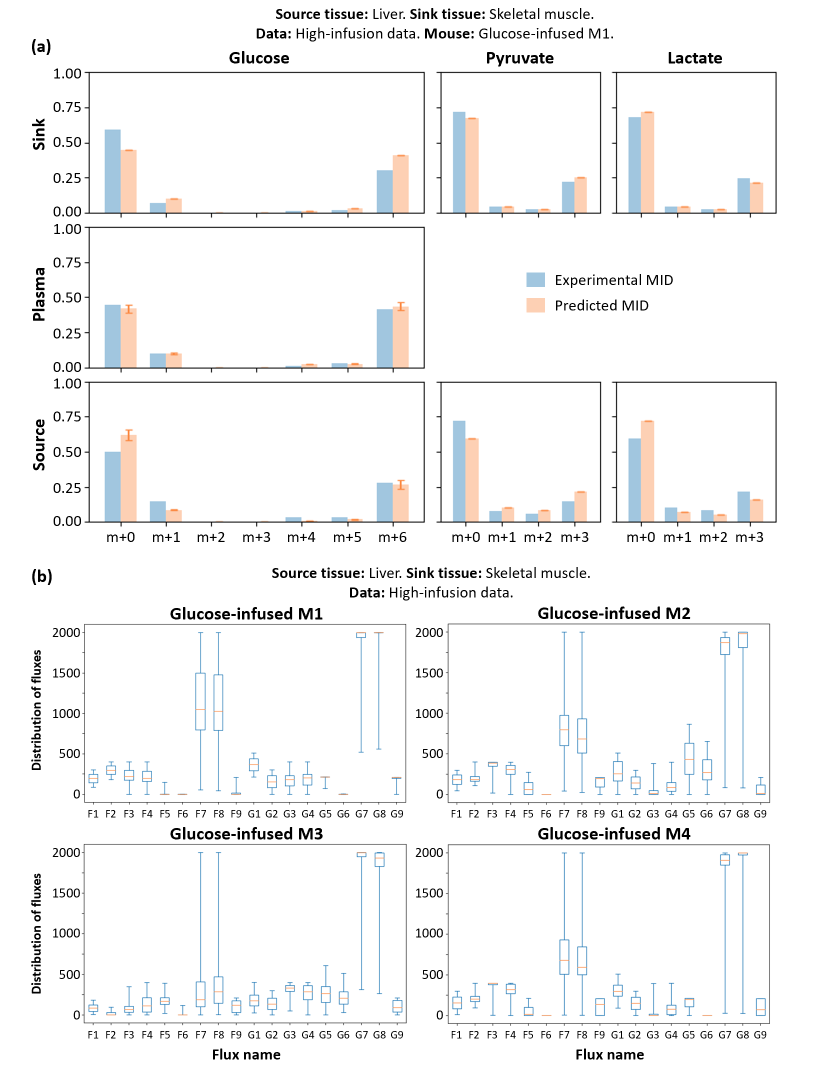

**Schematic 12**. Detailed information for the high-infusion system. (a) Comparison of experimental and predicted MID in the model. Average of predicted MID in all feasible solutions are displayed. Standard deviation is also displayed as error bar. Experimental MID could be well predicted by this model. Notably, the ratio of high 13C isotopomer is significantly higher than that of the original system. (b) Distribution of 18 variable fluxes in all feasible solutions. (c) Distribution of cost function fitted with data from different mice or unfitted control data. U-statistics of rank-sum test and p-values are displayed. Distribution of glucose contribution shows glucose contributes more than lactate to the TCA cycle. All subfigures are fitted with different glucose-infused mice from the high-infusion data. Source tissue is liver and sink tissue is skeletal muscle. Specifically, subfigure (a) is based on data from glucose-infused mouse M1. In all box plots, boxes represent quantiles and whiskers represent extremes.

The definition of glucose contribution in the complete model $R_{glc}^{'}$ is also same (figure S4C). Distribution of $R_{glc}^{'}$ in unfiltered results is slightly different from filtered ones (Schematic 12D, compared with figure S4D).

#### Model C: model for three tissues

| Category | Parameter | Comment | Value |
| --- | --- | --- | --- |
| Model |  | Maximal flux value | 1000 |
|  |  | Value of supplement glucose flux in source tissue | 100 |

Similarly, the fitting process except filtering is identical to filtered results of Model C. MID prediction and distribution of objective function in unfiltered results are almost same as filtered results (Schematic 13A, C, figure S5A, S5C). The distribution of fluxes in feasible solutions has a significant change: because the is higher in this calculation, H7 and H8 are slightly larger than other fluxes (Schematic 13B, compared with figure S5B). In this model, the filter just requires G9 and H9 not to be too small at the same time. Therefore, in unfiltered results, F9 and G9 tend to be optimized to extreme value (Schematic 13B), while in filtered ones F9 tends to concentrate to a small value, and G9 tends to concentrate to a large value (figure S5B).

The definition of glucose contribution in the complete model $R_{glc}^{'}$ is also same (figure S5D). Distribution of $R_{glc}^{'}$ in unfiltered results is similar but also slightly different from filtered ones (Schematic 13D, compared with figure S5E).

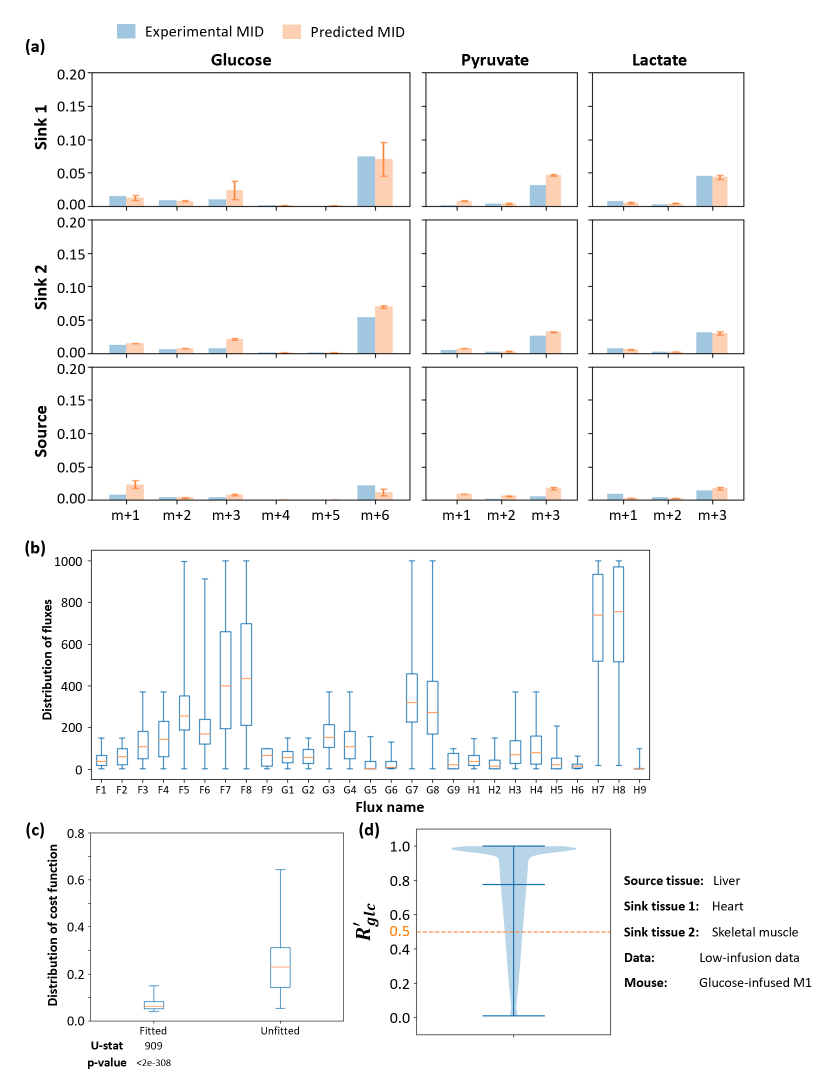

**Schematic 13**. Detailed information of the multi-tissue model. (a) Comparison of experimental and predicted MID in the model. Average of predicted MID in all feasible solutions are displayed. Standard deviation is also displayed as error bar. Experimental MID could be well predicted by this model. Because of the low abundance, only isotopomers with more than one 13C are displayed. (b) Distribution of 27 variable fluxes in all feasible solutions. (c) Distribution of cost functions of feasible solutions compared with that from unfitted control. U-statistics of rank-sum test and p-values are displayed. (d) Distribution of glucose contribution shows glucose contributes more than lactate to the TCA cycle. In all subfigures, the model is fitted with glucose-infused mouse M1 from the low-infusion data in Hui et al, 2017. The source tissue is liver and the sink tissue 1 and 2 are heart and skeletal muscle respectively. In all box plots, boxes represent quantiles and whiskers represent extremes.

#### Model D: model for three circulating metabolites for low-infusion data

| Category | Parameter | Comment | Value |
| --- | --- | --- | --- |
| Model |  | Maximal flux value | 1000 |
|  |  | Value of supplement glucose flux in source tissue | 200 |

#### Model E: model for three circulating metabolites for high-infusion data

| Category | Parameter | Comment | Value |
| --- | --- | --- | --- |
| Model |  | Maximal flux value | 2000 |
|  |  | Value of supplement glucose flux in source tissue | 150 |

The fitting process in this two calculations except filtering are identical to filtered results of Model D and E. MID prediction and distribution of objective function in unfiltered results are almost same as filtered results (Schematic 14A-B, E-F, figure S6A-B, E-F). Due to higher value of , range of all fluxes are larger in this calculation than that in main figures, especially for G7 and G8 in Model E (Schematic 14D, compared with figure S6D). Filtering also dramatically change distributions of TCA fluxes F11 and G11 in two models: in unfiltered results, F11 concentrates on maximum value while G11 concentrates on minimum value in Model D (Schematic 14C), and in Model E their quantiles are also closed to extreme value (Schematic 14D). However, in filtered results F11 and G11 concentrate on intermediate values in both Model D and Model E (figure S6C-D).

The definition of contribution from different metabolites in the complete model $R_{glc}^{'}$, $R_{lac}^{'}$ and $R_{pyr}^{'}$ are also same (figure S6H). Compared with filtered results, unfiltered results shows almost zero $R_{pyr}^{'}$ and lower $R_{lac}^{'}$, as well as relatively higher $R_{glc}^{'}$ (Schematic 14G-H, compared with figure S6I-J). This may be due to much higher median value of F4 and F10 in this unfiltered results compared to filtered results (Schematic 14C, figure S6C). Considering that the function of source tissue (liver) is converting lactate and other carbon source to glucose to supply other organs, lower outflux of lactate (F4) and pyruvate (F10) in source tissue in filtered results are more physiologically feasible.

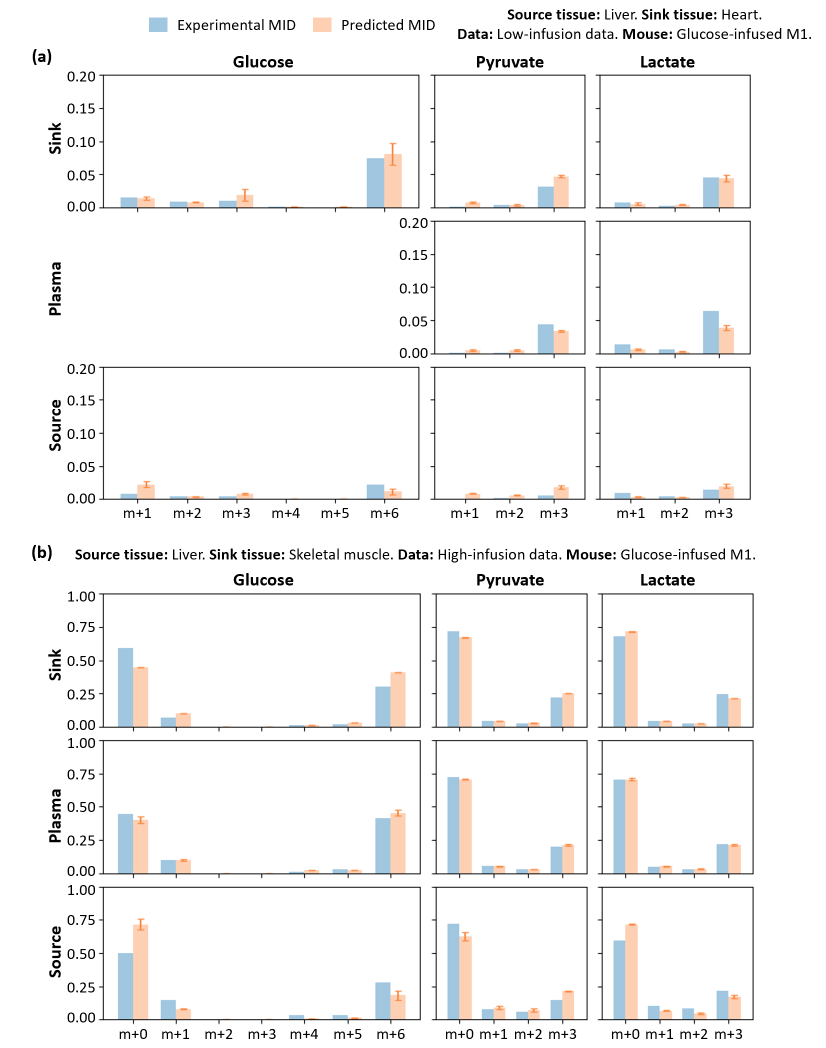

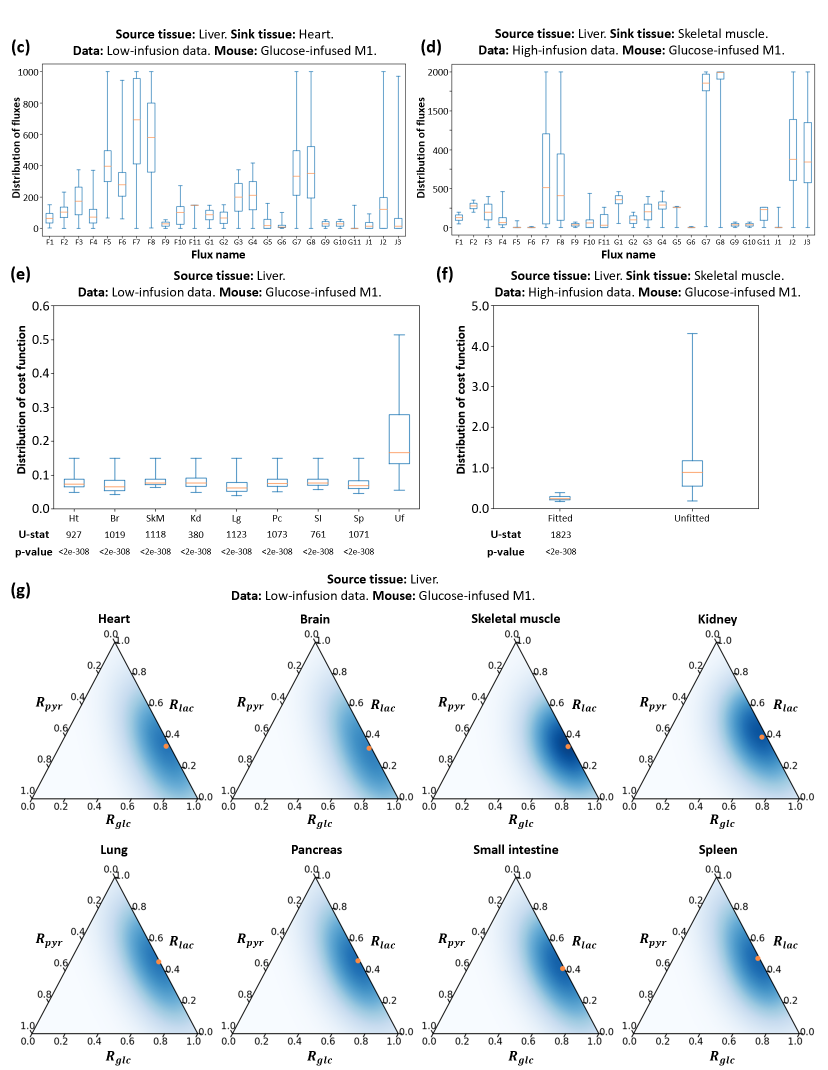

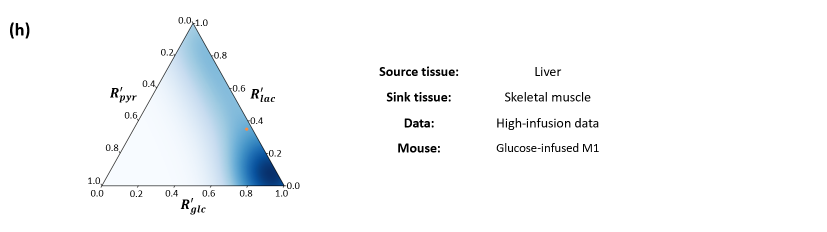

**Schematic 14**. Detailed information of the model with multiple circulating metabolites. (a, b) Comparison of experimental and predicted MID in the model fitted with the low-infusion data (a) or the high-infusion data (b). Average of predicted MID in all feasible solutions are displayed. Standard deviation is also displayed as error bar. Experimental MID could be well predicted by this model. (c, d) Distribution of 25 variable fluxes in all feasible solutions fitted with the low-infusion data (c) or the high-infusion data (d). (e, f) Distribution of cost functions of feasible solutions compared with that from random unfitted control, fitted with the low-infusion data (e) or the high-infusion data (f). U-statistics of rank-sum test and p-values are displayed. (g, h) Contributions to the TCA cycle when additional nutrients are considered, fitted with the low-infusion data (g) or the high-infusion data (h). For most sink tissues, glucose contributes to the TCA cycle more than lactate and pyruvate, especially in heart, brain, skeletal muscle, kidney and small intestine. The orange point indicates average level.

Subfigure (a), (c), (e) and (g) are fitted with glucose-infused mouse M1 from the low-infusion data in Hui et al, 2017, and source tissue is liver. Specifically, sink tissue in (a) and (c) is heart. Subfigure (b), (d), (f) and (h) are fitted with glucose-infused mouse M1 from the high-infusion data, and the sink tissue and source tissue are liver and skeletal muscle respectively.
